# Mitochondrial inner membrane partners of Aurora kinase A/AURKA and PHB2 shape organelle metabolic heterogeneity

**DOI:** 10.64898/2026.03.17.712317

**Authors:** Claire Caron, Nicolas Y. Jolivet, Diala Kantar, Étienne Coyaud, Giulia Bertolin

## Abstract

Mitochondria support tumor growth by flexibly rewiring their own activity. Yet, how signaling networks organize this metabolic diversity across and within cells remains poorly understood. Here, we uncover a platform of inner mitochondrial membrane proteins that couple Aurora kinase A (AURKA) and the mitophagy receptor Prohibitin-2 (PHB2) to the local control of ATP production and organelle architecture. Using proximity interactomics and live-cell FRET/FLIM microscopy, we identify NDUFA9, ATP5F1A/B, SAMM50, and SLC25A13 as shared AURKA/PHB2 interactors positioned at respiratory chain and *cristae*-maintaining sites. We show that AURKA overexpression profoundly rewires the interactomes of NDUFA9 and ATP5F1A without disrupting their nanoscale proximity in the inner membrane, and reshapes mitochondrial morphology while preserving these signaling hubs. The AURKA/PHB2-targeting compound HMBB restores NDUFA9 and ATP5F1A interactomes, indicating that these complexes act as key nodes for AURKA-dependent metabolic adaptation at the population level. Finally, single-cell FRET microscopy coupled to super-resolution imaging reveals that SLC25A13 is required to sustain AURKA-induced metabolic heterogeneity within individual cancer cells. Our work links a multifunctional kinase, a mitophagy receptor, and respiratory complexes into a common inner membrane platform that spatially drives mitochondrial heterogeneity, with implications for metabolism-focused anticancer strategies.

## Introduction

Mitochondria are a pivotal source of energy for a variety of cancers, including breast malignancies (Whitaker-Menezes *et al*., 2011; Sotgia *et al*., 2012). These organelles provide energy through oxidative phosphorylation (OXPHOS).

In cancer, it appears increasingly clear that metabolism can be reprogrammed in a cell-autonomous manner. This leads to the appearance of heterogeneous metabolic phenotypes that contribute to disease onset and evolution (Kim & DeBerardinis, 2019). Efforts have been made to develop single-cell approaches capable of revealing and tracking metabolic heterogeneity (Evers *et al*., 2019). Advances in super-resolution imaging also showed that mitochondria are a structurally and functionally heterogeneous population in cells that heavily rely on OXPHOS.

Organelles devoid of *cristae* were shown to be present and metabolically active, sustaining the functioning of mitochondria in OXPHOS-active cells through the reductive pathway (Ryu *et al*., 2024). These recent results demonstrate that mitochondria are metabolically heterogeneous not only at the tissue or cell population levels, but also within individual cells.

The cell cycle-related protein AURKA is often overexpressed in breast and hematopoietic tumors (Nikonova *et al*., 2013), and regulates mitochondrial metabolism in OXPHOS-reliant cells (Sharma *et al*., 2023). This is made possible by the localization of the kinase within mitochondria, where it interacts with a variety of metabolism-related partners (Bertolin *et al*., 2018). When the kinase is overexpressed, it triggers mitophagy by interacting with the autophagy mitochondrial receptor Prohibitin-2 (PHB2) (Bertolin *et al*., 2021). This event induces the degradation of a portion of the mitochondrial network, while maintaining the organelles with high energetic capabilities (Bertolin *et al*., 2018). AURKA was shown to be in proximity with the mitochondrial Complex V subunits ATP5F1A and ATP5F1B (Sharma *et al*., 2023), but it remains to be established whether this functional interaction directly triggers the modulation of OXPHOS activity when the kinase is overexpressed. The recent combination of the genetically encoded Förster’s Resonance Energy Transfer (FRET) mitoGO-ATeam2 biosensor (Nakano *et al*., 2011) with Enhanced Super-Resolution Radial Fluctuations (SRRF) microscopy (Laine *et al*., 2023) provided a pipeline to experimentally detect metabolic heterogeneity in individual cells (Jolivet *et al*., 2026b). This approach revealed that transient AURKA overexpression dramatically increases metabolic heterogeneity in MCF7 breast cancer cells, with a large proportion of the kinase being activated directly on ATP5F1A-rich clusters. Although the pharmacological inhibition of AURKA lowered the degree of metabolic heterogeneity, the requirement of known partners of the kinase participating in this phenotype remains to be explored.

Here, we combine interactomics and FRET microscopy to show that AURKA and PHB2 associate with a set of Inner Mitochondrial Membrane (IMM) interactors. Among these, NDUFA9, ATP5F1A and B, SAMM50, and SLC25A13 show the strongest association profiles with both AURKA and PHB2. These interactors have established roles in the maintenance of mitochondrial metabolism and were recently identified as *cristae*-maintaining proteins (Schaumkessel *et al*., 2025). We also show that AURKA overexpression induces profound rewiring of the NDUFA9 and ATP5F1A interactomes. The newly developed AURKA/PHB2-targeting compound HMBB (Djehal *et al*., 2026) restores the interactomes of NDUFA9 and ATP5F1A, suggesting that these two partners are key platforms for AURKA-dependent metabolic adaptation at the cell-population level. This is also true at the single-cell level, where the combination of FRET and SRRF microscopy reveals that SLC25A13 maintains metabolic heterogeneity in AURKA-overexpressing cells. Altogether, we identify a platform of AURKA/PHB2-associating IMM proteins that governs spatial and functional mitochondrial heterogeneity when AURKA is overexpressed.

## Results

### AURKA and PHB2 co-interact with mitochondrial inner membrane proteins

We have previously reported the functional interaction between AURKA and the Prohibitin (PHB) complex (Bertolin *et al*., 2021; Djehal *et al*., 2026). Within the PHB complex, PHB2 is required for AURKA to trigger mitochondrial clearance by mitophagy. Along with a significant portion of the mitochondrial network being lost, the remaining organelles show dramatic reorganization of their ultrastructure and overall network morphology (Bertolin et al., 2018, 2021; Djehal et al., 2026). Since these events are AURKA- and PHB2-dependent, we reasoned that these proteins require a common set of functional interactors to regulate such a diversity of organelle functions. To investigate this, we generated the proximity interactome of PHB2 using BioID approaches (Roux et al., 2012). We used HEK293 Flp-In T-REx cells that stably express PHB2 fused to a mutated biotin ligase (BirA*) tag at the C-terminus. Tetracycline was used to induce the expression of PHB2-BirA*, and biotin was then used to label proximal proteins in a 10-20 nm range from PHB2 in living cells (Table 1, Supplementary Fig. 1).

After removing the background noise from the PHB2 interactome, we selected mitochondrial proteins with a *log2* fold change ≥1 and a *q* value < 0.01 (Fig. 1A). Fifty-three proteins were considered as PHB2 high-confidence proximal interactors, and they were then intersected with 147 mitochondrial hits from a previously-generated interactome of AURKA (Bertolin et al., 2018) (Fig. 1A). This analysis led to the identification of 21 mitochondrial proteins shared between the PHB2 and AURKA interactomes. Gene Ontology analyses showed that several hits are localized at the IMM, in proximity to the respiratory chain (Vercellino & Sazanov, 2022). Functionally, these proteins participate in mitochondrial-specific molecular pathways, such as oxidative phosphorylation, ATP production, and metabolism (Fig. 1C).

**Fig. 1.**
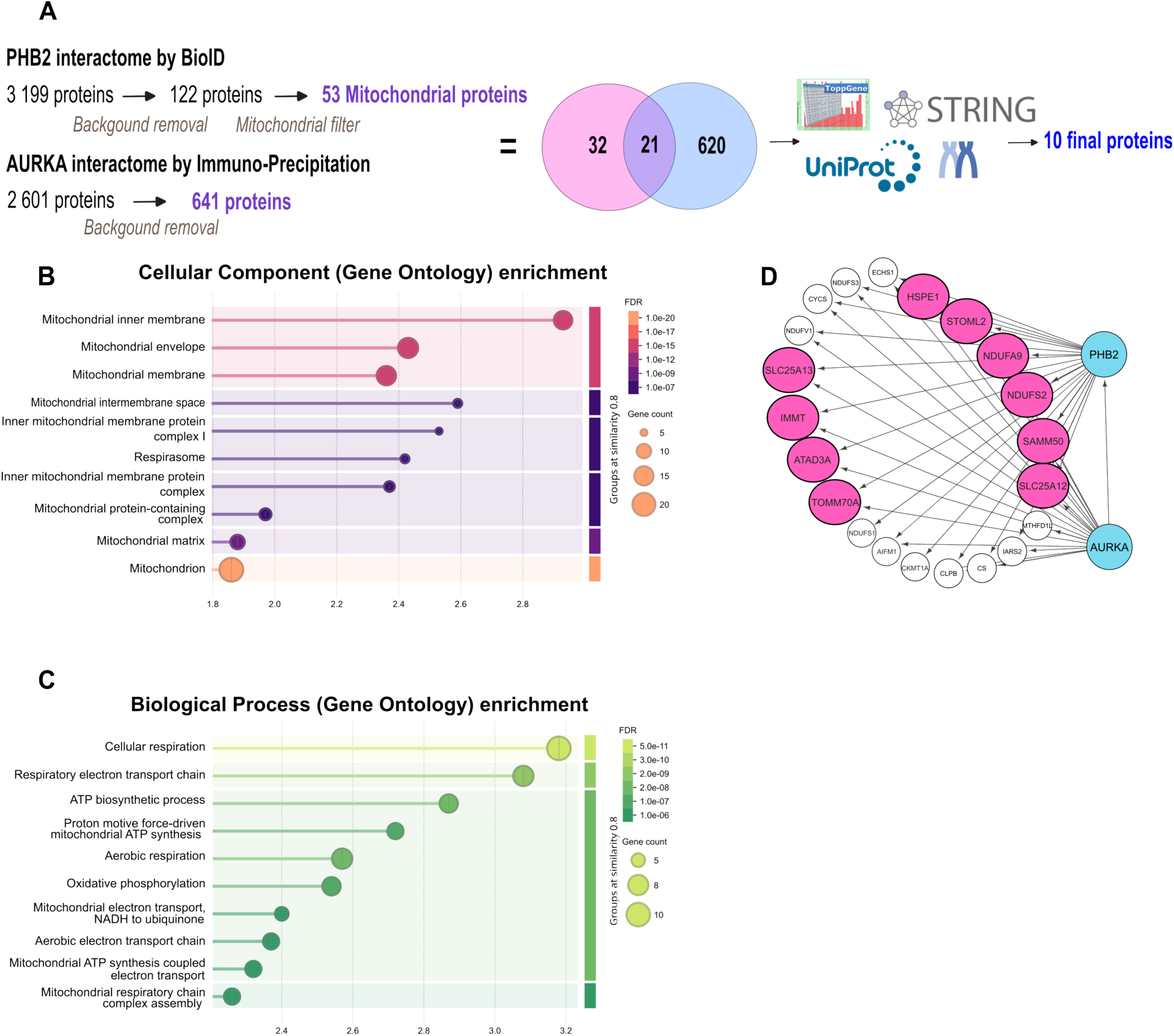
The AURKA/PHB2 cross-interactome identifies common proximal interactors at the IMM. (**A**) Comparison between the list of AURKA interactors obtained by immunoprecipitation and the list of PHB2 interactors obtained by BioID. The Venn diagram represents the common interactors. The final selection of the 10 most abundant interactors of AURKA and PHB2 was made based on enrichment analyses and annotations obtained from ToppGene, Metascape, UniProt, and the STRING databases. (**B-C**) Functional enrichment analysis of the mitochondrial protein set using the STRING database. The x-axis indicates the enrichment score. The size of the dots corresponds to the number of proteins of each given category. The color of the dots represents the false positive rate (FDR) as indicated. Terms are grouped by similarity (threshold = 0.8), grouping functionally related annotations with gene sets that overlap by >80%. (**D**) Cytoscape representation of the 21 mitochondrial proteins common to AURKA and PHB2. The 10 selected proteins are shown in pink.

Since AURKA and PHB2 functionally interact on the IMM (Durel et al., 2021; Bertolin et al., 2021; Djehal et al., 2026), we selected proteins known to be located exclusively at this location or on contact sites between the IMM and the Outer Mitochondrial Membrane (OMM), and with the highest *log2* fold change among the cross-interactome hits (Fig. 1A). Overall, we identified 10 AURKA/PHB2 shared hits for comparative analyses, *i.e.*, HSPE1, STOML2, NDUFA9, NDUFS2, SAMM50, SLC25A12, SLC25A13, IMMT/MIC60, ATAD3A, and TOMM70 (Fig. 1A, D).

Förster’s Resonance Energy Transfer/Fluorescence Lifetime Imaging Microscopy (FRET/FLIM) was used to validate the protein-protein vicinities in live MCF7 cells. AURKA and PHB2 were fused to the FRET donor GFP, while each of the 10 selected partners was fused to the acceptor fluorophore mCherry. FRET/FLIM analyses revealed that AURKA and PHB2 show a significant proximity with the Complex I subunits NDUFA9 and NDUFS2, the OMM/IMM contact site and import-related protein SAMM50, and with the aspartate/glutamate antiporter SLC25A13 (Fig. 2A-C, Supplementary Fig. 2, 3). Since the proximity between AURKA and the mitochondrial Complex V has already been described (Sharma *et al*., 2023), we used it as a positive control for FRET/FLIM analyses (Fig. 2A). Indeed, we retrieved the proximity between AURKA and the Complex V subunits ATP5F1A and ATP5F1B (Fig. 2A, Supplementary Fig. 2). We also revealed the proximity between PHB2 and both Complex V subunits (Fig. 2B, Supplementary Fig. 3). This demonstrates that Complex V is in proximity to both moieties of the AURKA/PHB2 complex. Furthermore, we confirmed the proximity between AURKA and TOMM70A (Bertolin *et al*., 2018), although this subunit of the TOMM import complex was not found to interact with PHB2 in FRET/FLIM analyses (Supplementary Fig. 2, 3). The MICOS subunit IMMT/MIC60 showed a significant proximity with PHB2, but not with AURKA (Supplementary Fig. 2, 3), whereas the mitochondrial *cristae* protein ATAD3A showed a significant, albeit variable proximity with AURKA and PHB2 (Supplementary Fig. 2, 3). Finally, the cardiolipin-related protein STOML2 was excluded from subsequent analyses because its ectopic expression led to mislocalization in the nucleus (Supplementary Fig. 2), This was also the case for the mitochondrial co-chaperonin CH10/HSPE1, which showed an intense cytoplasmic signal upon expression (Supplementary Fig. 2, 3).

**Fig. 2.**
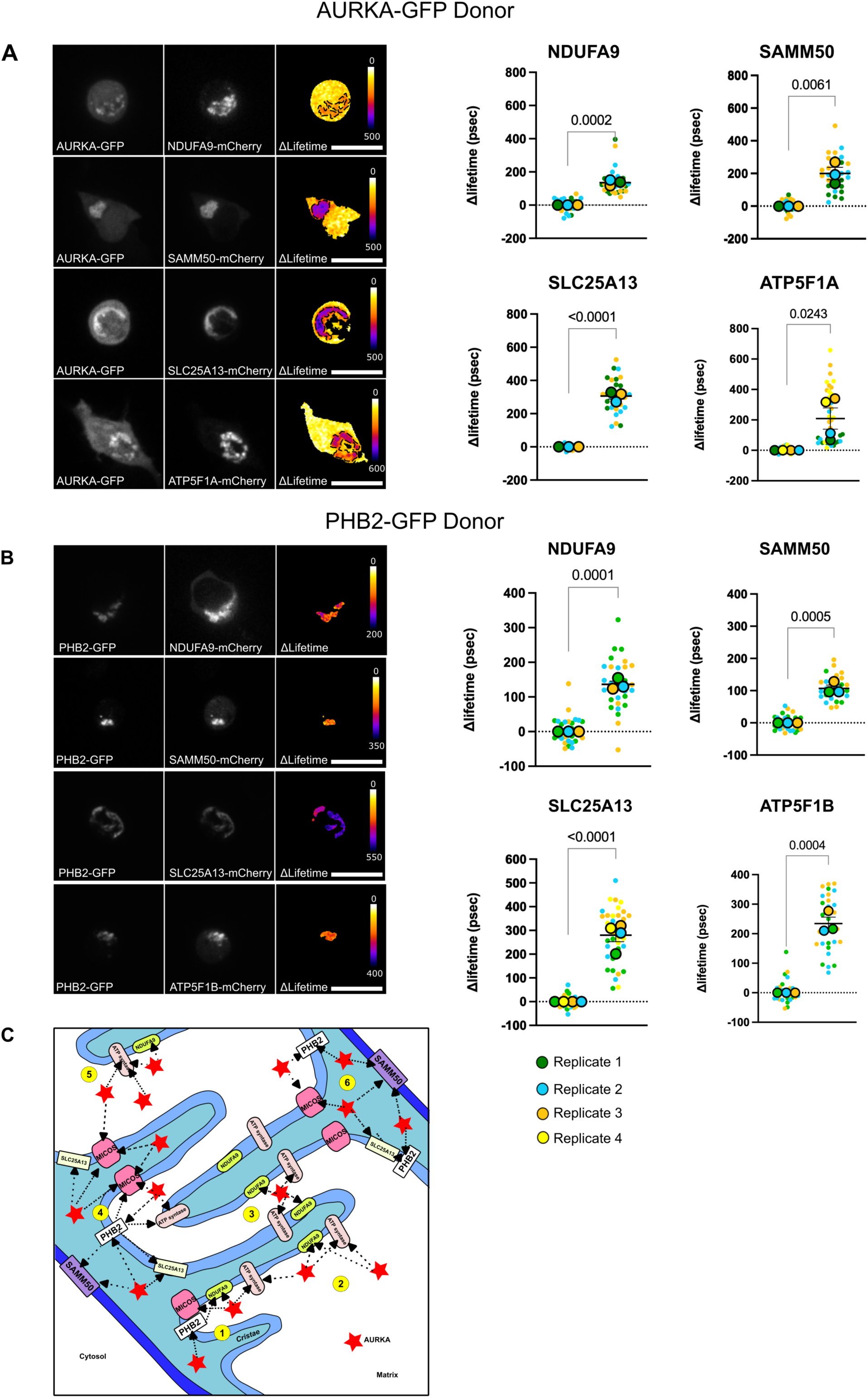
(Figure on the previous page) AURKA and PHB2 co-interact with NDUFAG, SAMM50, SLC25A13, and mitochondrial Complex V subunits. **(A-B)** Fluorescence, FRET/FLIM micrographs (left) and corresponding quantitative analyses (right) of MCF7 cells co-expressing either AURKA-GFP (**A**) or PHB2-GFP (**B**) (donors), and each of the indicated IMM proteins fused to mCherry (acceptors). Pseudocolor scale: pixel-by-pixel ΔLifetime. Dotted area: acceptor-rich mitochondrial area. Scale bars: 5 µm. *n* = 10 cells per condition (small symbols) in each of three or four biological replicates. Large symbols indicate mean values for each biological replicate. Data are means ± S.D. Exact *P*-values are indicated for each comparison. (**C**) Cartoon representing all potential proximity scenarios (1-5) of AURKA (red star) with the IMM proteins PHB2, SAMM50, SLC25A13, the ATP synthase, NDUFA9 and the MICOS complex.

Overall, orthogonal interactomics and FRET/FLIM analyses revealed a shared set of interactors between AURKA and PHB2, located at the IMM or involved in IMM/OMM contact sites. These include subunits of Complex I such as NDUFA9, SAMM50, and SLC25A13.

### AURKA-dependent alterations of mitochondrial morphology do not modify protein proximities at the IMM

After detecting the proximity of both AURKA and PHB2 with IMM proteins, we sought to determine whether this proximity is driven by changes in mitochondrial ultrastructure. Previous reports showed that the overexpression of AURKA induces dramatic changes in mitochondrial morphodynamics, with the presence of swollen and aggregated organelles (Grant *et al*., 2018; Bertolin *et al*., 2018). Therefore, we asked whether these morphological changes may induce or modify the proximity between AURKA or PHB2 and selected IMM proteins. To this end, we used direct stochastic optical reconstruction microscopy (dSTORM). dSTORM is a super-resolution microscopy approach providing an optical resolution of 10-20 nm (Heilemann *et al*., 2008), well suited to investigate mitochondrial architecture and composition (Jakobs, 2006; Huang *et al*., 2008; Dlasková *et al*., 2018).

We performed dSTORM imaging in MCF7 cells expressing an empty vector or AURKA-6xHis. These cells were co-labeled for AURKA (Fig. 3A) and MIC60, or for PHB2 and ATP5F1B (Fig. 3B). These proteins are widely used as *bona fide* markers of the IMM, with MIC60 detecting *cristae* junctions (CJ) and ATP5F1B detecting *cristae* membranes (CM) (Dlasková *et al*., 2019). Compared to cells transfected with a control vector, MIC60 and ATP5F1B showed organelle swelling and aggregation upon AURKA overexpression, as expected (Fig. 3A-B). We also detected the presence of both endogenous AURKA and AURKA-6xHis in the mitochondrial matrix (Fig. 3A), expanding on a previous report (Durel *et al*., 2021). Quantitative colocalization analyses using the GCoPS software (Lavancier *et al*., 2020; Durel *et al*., 2021) revealed that the AURKA/MIC60 molecular proximities were identical in control cells and in cells overexpressing AURKA. A similar behavior was observed for the PHB2/ATP5F1B proximities (Fig. 3C).

**Fig. 3.**
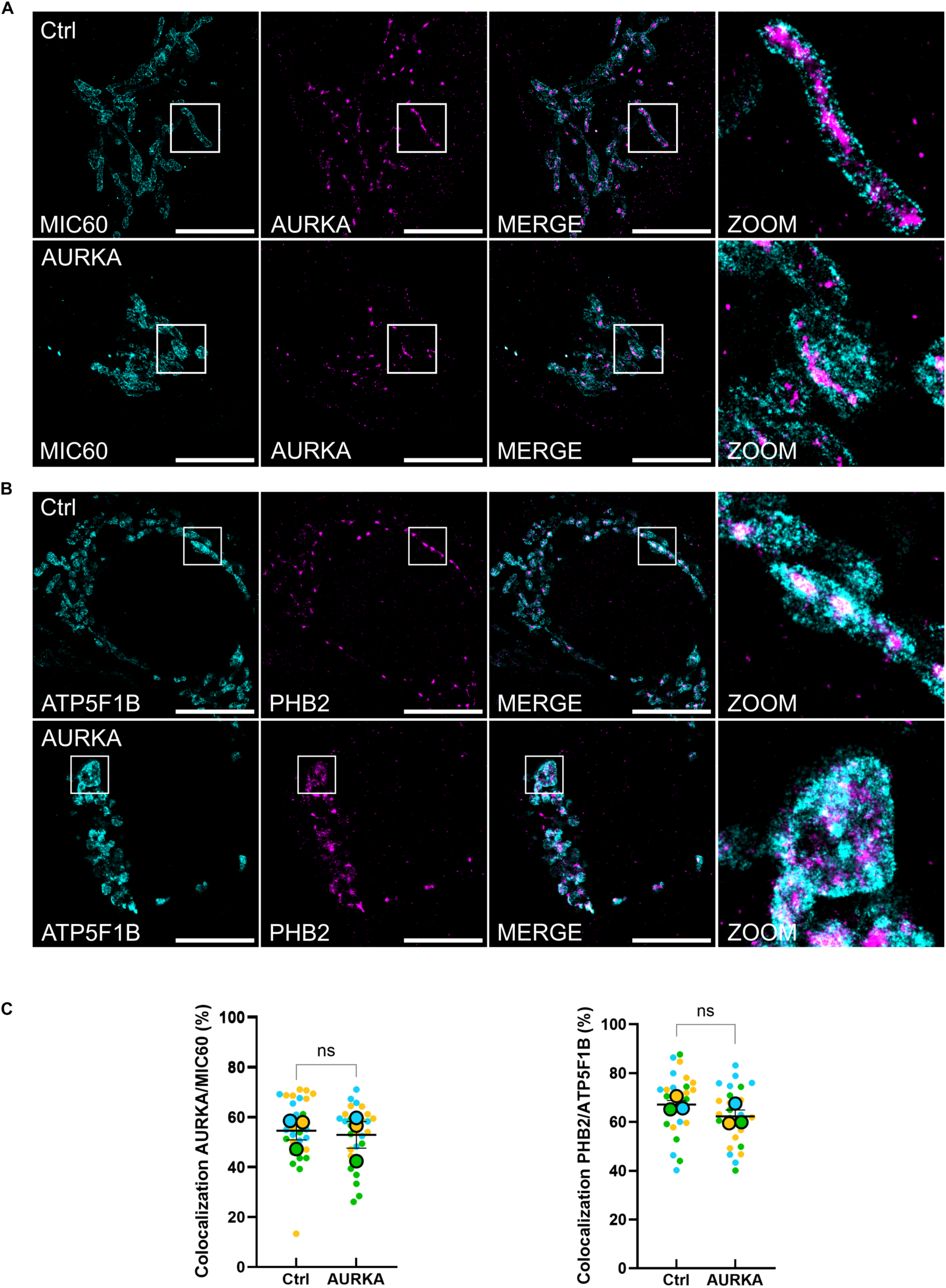
AURKA overexpression does not change protein proximities between IMM canonical markers. Maximal projections of representative 2D dSTORM super-resolution micrographs from MCF7 transfected with an empty vector (Ctrl) or AURKA-6XHis (AURKA), and co-stained for AURKA and MIC60 (**A**), or for ATP5F1B and PHB2 (**B**). The anti-MIC60 and ATP5F1B primary antibodies were detected using an Alexa 647-conjugated secondary antibody and pseudocolored cyan, while anti-AURKA and PHB2 primary antibodies were detected using a CF532 secondary antibody and pseudocolored magenta. Insets: magnified region where each staining is merged (ZOOM). Scale bar: 10 *μ*m. (C) Colocalization analyses performed with the GCoPS software, showing the percentage of colocalization for each protein pair in controls and AURKA-overexpressing cells. *n* = 10 cells per condition (small symbols) in each of three biological replicates. Large symbols indicate mean values for each biological replicate. Data are means ± S.D. ns: not significant.

Therefore, the overexpression of AURKA induces a reorganization of the mitochondrial network and changes in mitochondrial morphology. However, molecular proximities involving AURKA or PHB2 with CJ and CM markers are not modified by these morphological alterations.

### AURKA overexpression rewires the molecular interactomes of NDUFA9 and ATP5F1A

Previous reports showed that AURKA overexpression alters mitochondrial morphology and turnover rates to enhance the metabolic capacity of the mitochondrial network, and this in a PHB2-dependent manner (Bertolin *et al*., 2018, 2021; Sharma *et al*., 2023). However, the molecular partners acting with AURKA and PHB2 in the regulation of mitochondrial ATP rates remain to be fully elucidated. The Complex V subunits ATP5F1A and ATP5F1B have previously been identified as metabolism-related interactors of AURKA (Sharma *et al*., 2023). In addition to AURKA, FRET/FLIM analyses revealed that these proteins are also interactors of PHB2, along with Complex I subunits (Fig. 2 and Supplementary Fig. 2, 3). We therefore hypothesized that Complex I or Complex V subunits may be hubs of metabolic modulation in AURKA-overexpressing cells. If that were the case, AURKA overexpression may induce changes in Complex I and Complex V interactomes, which are indicative of a rewired mitochondrial metabolism.

To test this hypothesis, we chose NDUFA9 and ATP5F1A as significant co-interactors of AURKA and PHB2 (Fig. 2 and Supplementary Fig. 2, 3), and as proxies for Complex I and V, respectively. NDUFA9 and ATP5F1A were fused to TurboID tag, a catalytically faster mutant of BirA* (Branon *et al*., 2018), at the C-ter and stably inserted in HEK293 Flp-In T-REx cells overexpressing AURKA-6xHis or an empty vector. Cells were then treated with Tetracycline for NDUFA9- or ATP5F1A-TurboID fusion protein expression. After biotin addition, mitochondrial fractions were isolated and corresponding protein lysates subjected to biotin-streptavidin purification followed by mass spectrometry to retrieve the proximal interactome of both baits (Supplementary Fig. 4).

Mass spectrometry analyses in control cells revealed that NDUFA9 interactors are involved in the generation of precursor metabolites and energy, respiratory chain complex assembly, mitochondrial membrane organization, and mitophagy (Fig. 4A). Overexpression of AURKA did not lead to the appearance of new metabolism-related GO terms, but to a redistribution of proteins among the four GO classes identified. When analyzing the ATP5F1A interactome, mass spectrometry analyses in control cells revealed that the hits are mainly involved in the generation of precursor metabolites and energy, in the assembly of the respiratory chain, and in membrane organization (Fig. 4B). AURKA overexpression led to a diversification of active metabolic pathways, including the appearance of new proximities with partners involved in catabolism and valine metabolism. Under these conditions, the number of interactors involved in the assembly of the respiratory chain complexes increased as well (Fig. 4B). Notably, proteins regulating the stability, the assembly, and the functionality of Complex IV appeared, *e.g.* COA1 and SURF1, corroborating previous reports showing an upregulation of Complex IV protein abundance upon AURKA overexpression (Bertolin *et al*., 2018, 2021).

**Fig. 4.**
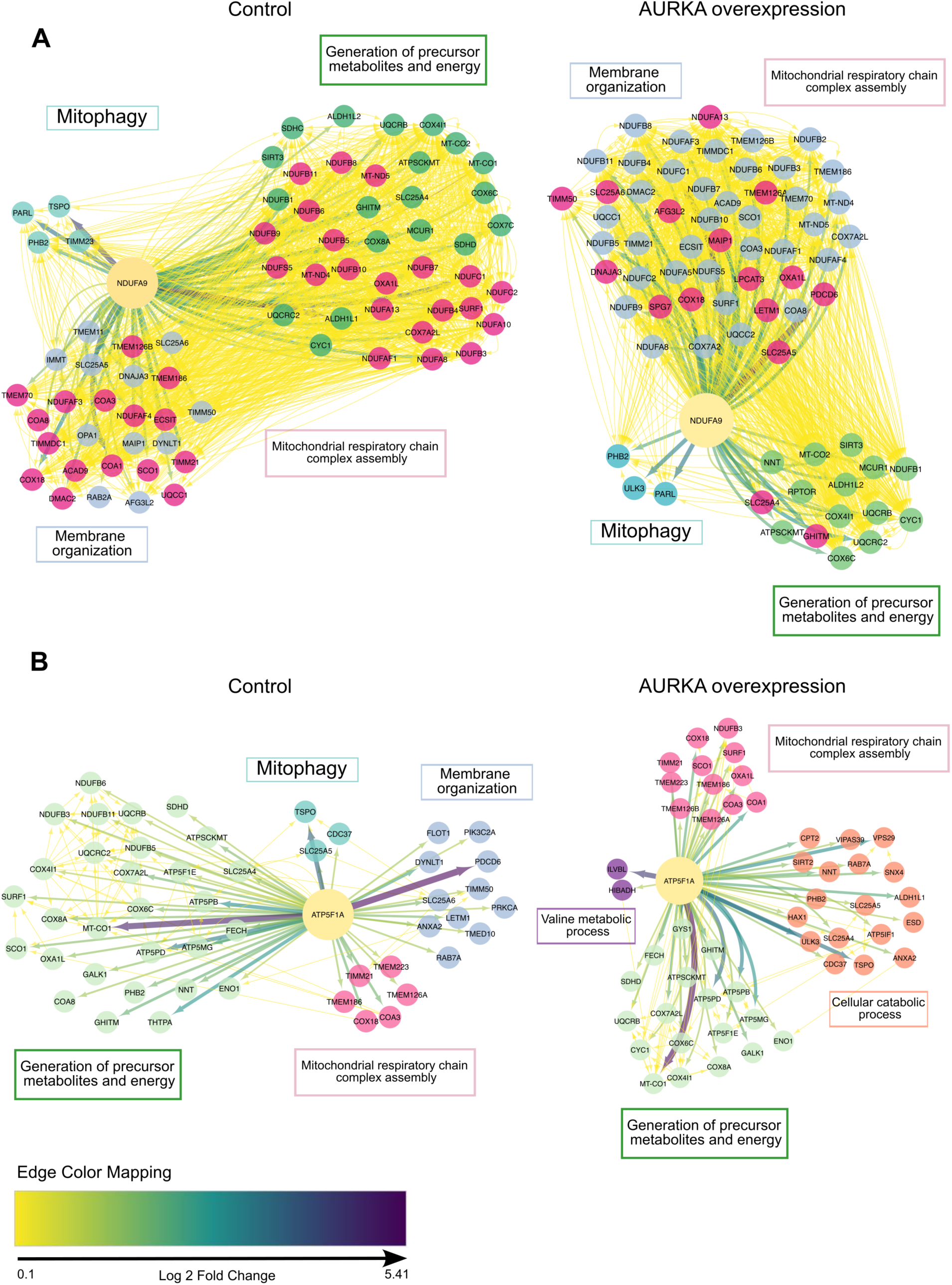
AURKA overexpression remodulates the NDUFAG and ATP5F1A proximal interactomes. (**A-B**) Cytoscape-based visualization of NDUFA9 (**A**) and ATP5F1A (**B**) proximal interactors. Hits were represented as follows: generation of precursor metabolites and energy class (GO: 0006091) in green; respiratory chain complex assembly (GO: 0033108) in pink; membrane organization (GO: 0061024) in grey; mitophagy (GO: 000423) in cyan; Valine metabolic process (GO: 0006573) in purple; cellular catabolic process (GO: 0044248) in orange. Arrows indicate an interaction. Arrow thickness: log2 fold change (graphically represented by a line thickness varying from 0 to 10 arbitrary units). Yellow depicts previously-reported protein-protein proximities, extracted from the STRIN9G database Version 12.0. For graphical purposes, the yellow edges thickness was arbitrarily set to a *|log2fc|=1*. Data are from three biological replicates.

In conclusion, AURKA overexpression leads to the diversification of metabolic networks by reshaping the mitochondrial interactome of ATP5F1A.

### The AURKA/PHB2-targeting compound HMBB restores Complex I and Complex V interactomes in AURKA-overexpressing cells

After observing that AURKA overexpression drives the onset of protein-protein proximities indicative of a rewired mitochondrial metabolism, we designed strategies to restore this phenotype. N-(4-hydroxy-3-methoxybenzyl)butyramide (HMBB) is a chemical compound that was recently characterized as a mitochondrial inhibitor of AURKA. By acting on the AURKA/PHB2 proximity, HMBB was shown to restore AURKA-related turnover by mitophagy (Djehal *et al*., 2026) and ATP production hotspots in mitochondria (Jolivet *et al*., 2026b).

We first explored the effect of HMBB on IMM protein proximities in T47D cells, which express high levels of endogenous AURKA (Bertolin *et al*., 2018; Sharma *et al*., 2023). T47D cells were treated with or without HMBB, co-stained for ATP5F1B and MIC60, and imaged using dSTORM super-resolution microscopy (Fig. 5). Employing this approach, we corroborated previous evidence showing the effect of HMBB on mitochondrial morphology and mass. Under these conditions, we observed that the mitochondrial network adopts a more heterogeneous morphology and that mass is partially increased (Djehal *et al*., 2026). However, GCoPS analyses revealed no differences in the MIC60/ATP5F1B colocalization levels in cells treated with vehicle or HMBB. Once more, these data indicate that changes in mitochondrial morphology do not drive profound alterations in IMM protein vicinities *per se*.

**Fig. 5.**
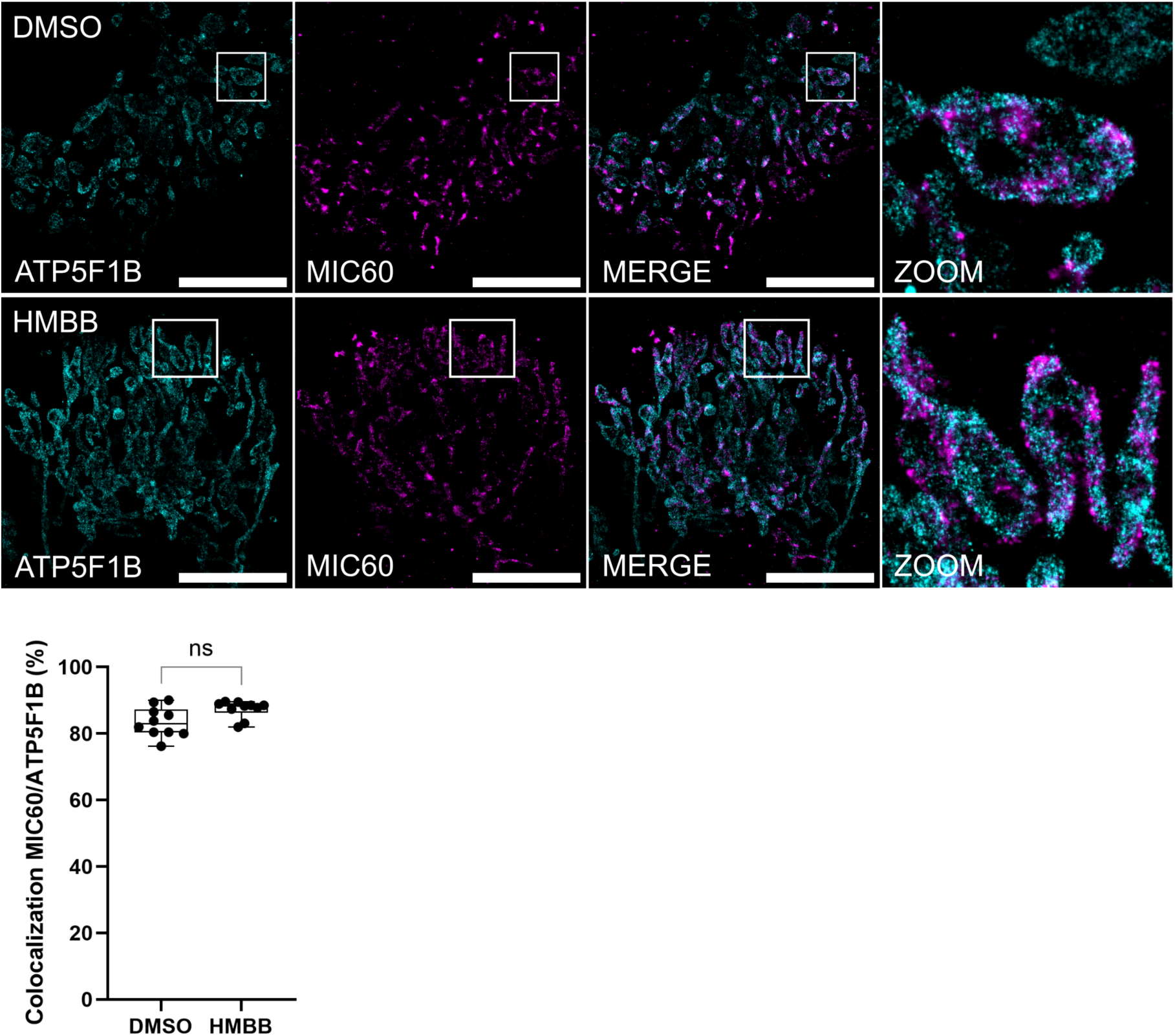
Treatment with HMBB does not change proximities between canonical IMM markers. (Upper panels) Maximal projections of representative 2D dSTORM super-resolution micrographs from T47D cells treated with DMSO or HMBB for 24 h, and co-stained for ATP5F1B and MIC60. The anti-ATP5F1B primary antibody was detected using an Alexa 647-conjugated secondary antibody and pseudocolored cyan, while the anti-MIC60 primary antibody was detected using a CF532 secondary antibody and pseudocolored magenta. Insets: magnified region where each staining is merged (ZOOM). Scale bar: 10 *μ*m. (Lower panel) Colocalization analyses performed with the GCoPS software, showing the percentage of colocalization between MIC60 and ATP5F1B in controls and HMBB-treated cells. *n* = 10 cells per condition in a representative biological replicate (of two). Data are means ± S.D. ns: not significant.

Next, we asked whether HMBB lowers mitochondrial ATP production by acting on the interactors of ATP5F1A and NDUFA9. These partners were chosen because of their direct involvement in OXPHOS composition and function. TurboID analyses were performed in mitochondrial fractions extracted from HEK293 Flp-In T-REx cells overexpressing AURKA-6xHis or an empty vector, and treated with vehicle or HMBB. Overexpression of AURKA in cells treated with DMSO increased the abundance of proximal interactors broadly involved in cellular metabolism and mitochondrial functions (*e.g.*, ALDH1L1, BLVRB, OPA3), with vesicular trafficking (*e.g.*, VIPAS39), and with the activation of translation (*e.g.*, EIF2B2) (Fig. 6A). HMBB treatment decreased the abundance of OPA3, BLVRB, VIPAS39, and EIF2B2. This suggests that HMBB could reverse protein-protein proximal interactions originally induced after AURKA overexpression. The effect of HMBB was even more striking when observing the NDUFA9 interactome. AURKA overexpression led to the loss of 11 interactors (Fig. 6B). No proximal interactor was seen as increased in abundance under these conditions. HMBB treatment not only restored but also increased the label-free quantification signals between NDUFA9 and seven out of the 11 interactors previously decreased in AURKA-overexpressing cells, including the IMM proteins COA1, TSPO, and ATAD3B. Furthermore, HMBB treatment enhanced the proximity detection signal between NDUFA9 and the IMM proteins OPA3, NUDT8, COX7C, and SDHAF4 (Fig. 6B).

**Fig. 6.**
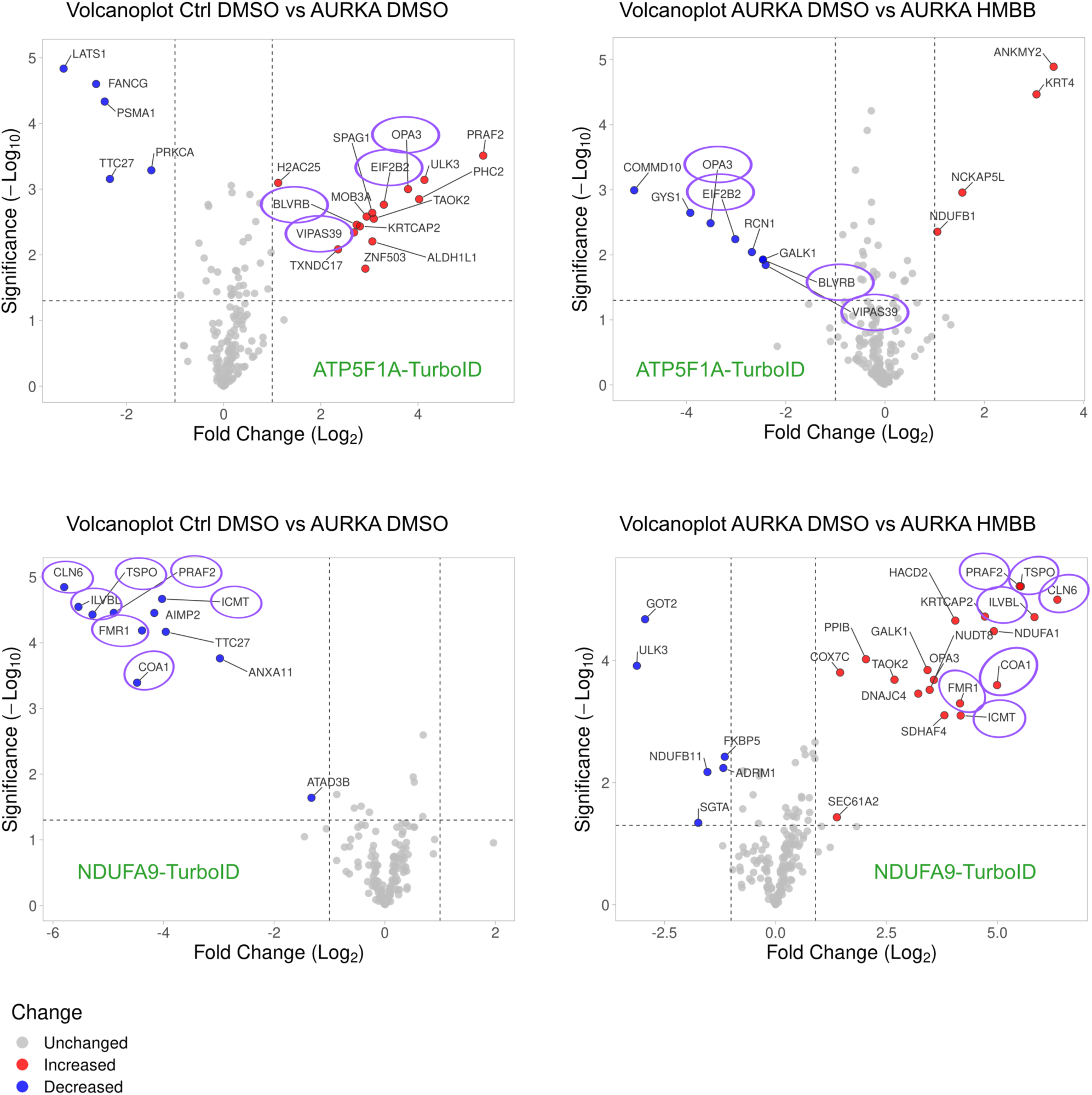
HMBB treatment restores the mitochondrial proximal interactome of NDUFAG and ATP5F1A. Proximal interactomics and TurboID pairwise comparisons from mitochondrial fractions of HEK293 Flp-In T-REx cells stably expressing ATP5F1A-TurboID (upper panels) or NDUFA9-TurboID (lower panels), transfected with an empty vector (Ctrl) or an AURKA-6XHis vector and treated with DMSO or HMBB for 6 h. Proteins with differential abundance were chosen by applying a *|log2fc|>1* and a *q*-value < 0.05. Proteins with increased abundance are indicated in red in the volcano plots, those with decreased abundance are indicated in blue, and those remaining unchanged are indicated in grey. Proteins labeled with a purple circle represent hits disappearing in each respective left panel and appearing in the corresponding right panel upon HMBB treatment. Data are from three biological replicates.

Overall, HMBB is an AURKA/PHB2-targeting compound restoring physiological protein-protein vicinities within mitochondrial Complex I and V, which are altered under AURKA overexpression conditions.

### SLC25A13 regulates PPARGC1 levels and mitochondrial ATP heterogeneity in AURKA-overexpressing cells

Upon the proximal interaction with the IMM proteins NDUFA9 and ATP5F1A, overexpressed AURKA modulates metabolism-related protein-protein vicinities. In parallel, AURKA has been shown to drive mitochondrial turnover by mitophagy through interacting with partners located at the same intramitochondrial location, such as PHB2 (Bertolin *et al*., 2021). Therefore, we asked whether AURKA and PHB2 co-interacting proteins could play a role in AURKA-dependent mitophagy.

We performed siRNA-mediated knockdown of *NDUFA9*, *SAMM50, SLC25A13,* and *ATP5F1B* in T47D cells and measured the relative area covered by the mitochondrial matrix marker PMPCB as a mitophagy readout (Djehal *et al*., 2026). While no significant variation was observed when NDUFA9, SAMM50, or ATP5F1B were depleted, the downregulation of *SLC25A13* led to a significant loss of PMPCB area normalized over the total cell area (Fig. 7A, Supplementary Fig. 5A). However, western blotting analyses revealed that the abundance of PMPCB was not significantly different in control and SLC25A13-depleted cells (Fig. 7B). This suggests that the apparent decrease in the relative mitochondrial area could be due to a change in mitochondrial morphology or organelle distribution within the cell. Mitochondrial length and branching were unaltered in the presence or absence of SLC25A13 (Supplementary Fig. 6A-C). In parallel, mitochondria in cells depleted for SLC25A13 are clustered around the perinuclear area, a phenotype not seen in control cells (Fig. 7A). It has been previously shown that mitochondrial clustering indicates lowered metabolic capacity (Wakim *et al*., 2017). Previous evidence also links enhanced breast cancer cell metabolism with high PPARGC1 (PGC1α+b) levels (Andrzejewski *et al*., 2017). Therefore, we tested whether mitochondrial clustering could be indicative of lowered PPARGC1 levels. Western blotting analyses revealed that PPARGC1A+B levels decrease in cells depleted for SLC25A13 (Fig. 7C).

**Fig. 7.**
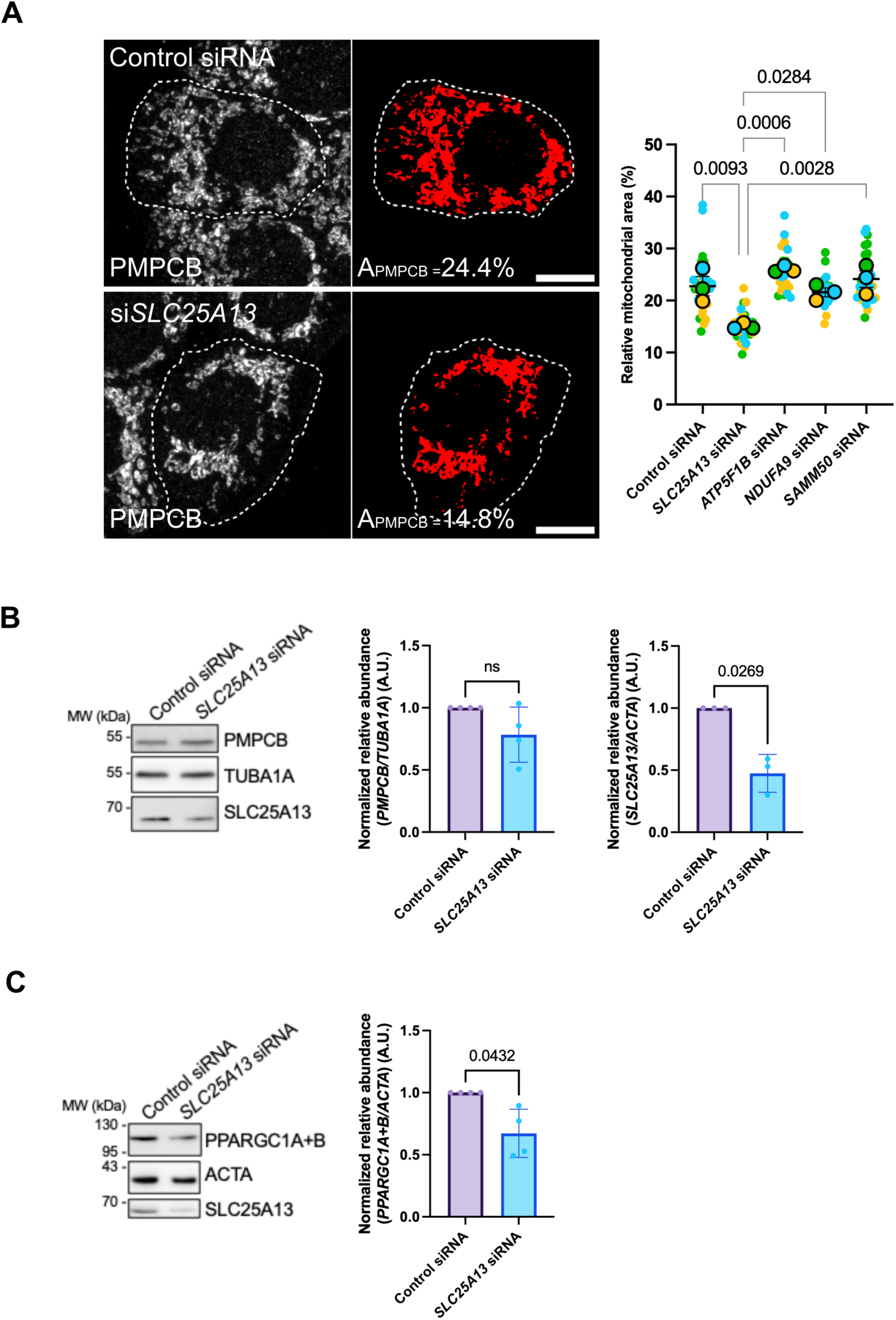
SLC25A13 depletion induces mitochondrial clustering and a decrease in PPARGC1 levels. (**A**) Relative mitochondrial area shown as the amount of PMPCB staining (threshold mask and corresponding quantification) in T47D cells transfected as indicated. A_PMPCB_: mitochondrial area normalized against total cell area (%). *n* = 10 cells per condition (small dots) in each of three biological replicates. Large dots indicate mean values for each replicate. Data are means ± S.D. Scale bar: 10 µm. (**B-C**) Representative western blot and corresponding quantifications of the relative abundance of PMPCB, SLC25A13 (**B**), and PPARGC1 (**C**) in total lysates of T47D cells transfected with a control or *SLC25A13*-specific siRNAs. Loading controls: TUBA1A (**B**), ACTA (**C**). Data are from *n* = 3 or 4 independent biological replicates, and presented as means ± S.D. Each dot corresponds to a biological replicate. A.U.: arbitrary units. Exact *P*-values are indicated for each comparison when significant.

SLC25A13 is an aspartate/glutamate transporter that plays key roles in energy metabolism (Tavoulari *et al*., 2022). To confirm that the decrease in PPARGC1A+B levels reflects alterations in ATP production, mitochondrial ATP production was measured by analyzing the FRET readout of the pH-insensitive mitoGO-ATEAM2 fluorescent biosensor (Nakano *et al*., 2011). First, mean ratiometric FRET index values confirmed that the depletion of SLC25A13 induces a significant decrease in mitochondrial ATP production (Fig. 8A). We then sought to understand whether this decrease in ATP production is evenly or heterogeneously distributed within the mitochondrial network. To this end, we used a previously established strategy combining FRET with Super Resolution Radial Fluctuations (SRRF) super-resolution microscopy to reveal ATP production *loci* with better spatial resolution than conventional techniques (Jolivet *et al*., 2026b). To identify ATP-producing or non-producing *loci*, FRET line analyses were performed. Fifty lines of 25 pixels each were randomly drawn on the entire mitochondrial network, and FRET index profiles were automatically calculated and extracted for each line, representing FRET variations along the line (Supplementary Fig. 7A). Lines with a Gaussian (FRET Hotspot) or inverse Gaussian (FRET Coldspot) profiles were extracted, representing FRET hotspots or coldspots, respectively. Then, the extent of FRET variations was used to infer the relative size of hotspots or coldspots. FRET variations were grouped into three clusters following silhouette coefficient and *k*-means analyses (Supplementary Fig. 7B, C), showcasing high, medium, or low FRET variations along the line. Given that lines have a constant size, the higher the FRET variation along the line, the smaller the ATP production site is. With this approach, previously termed *BioSenSRRF* (Jolivet *et al*., 2026b), ATP production *loci* can be visualized with superior spatial resolution, and the degree of metabolic heterogeneity measured with the mitoGO-ATeam probe can be determined. Of note, BioSenSRRF was shown to be compatible with different genetically-encoded FRET sensors, network morphologies and when mitochondria undergo different types of stress (Jolivet *et al*., 2026b).

**Fig. 8.**
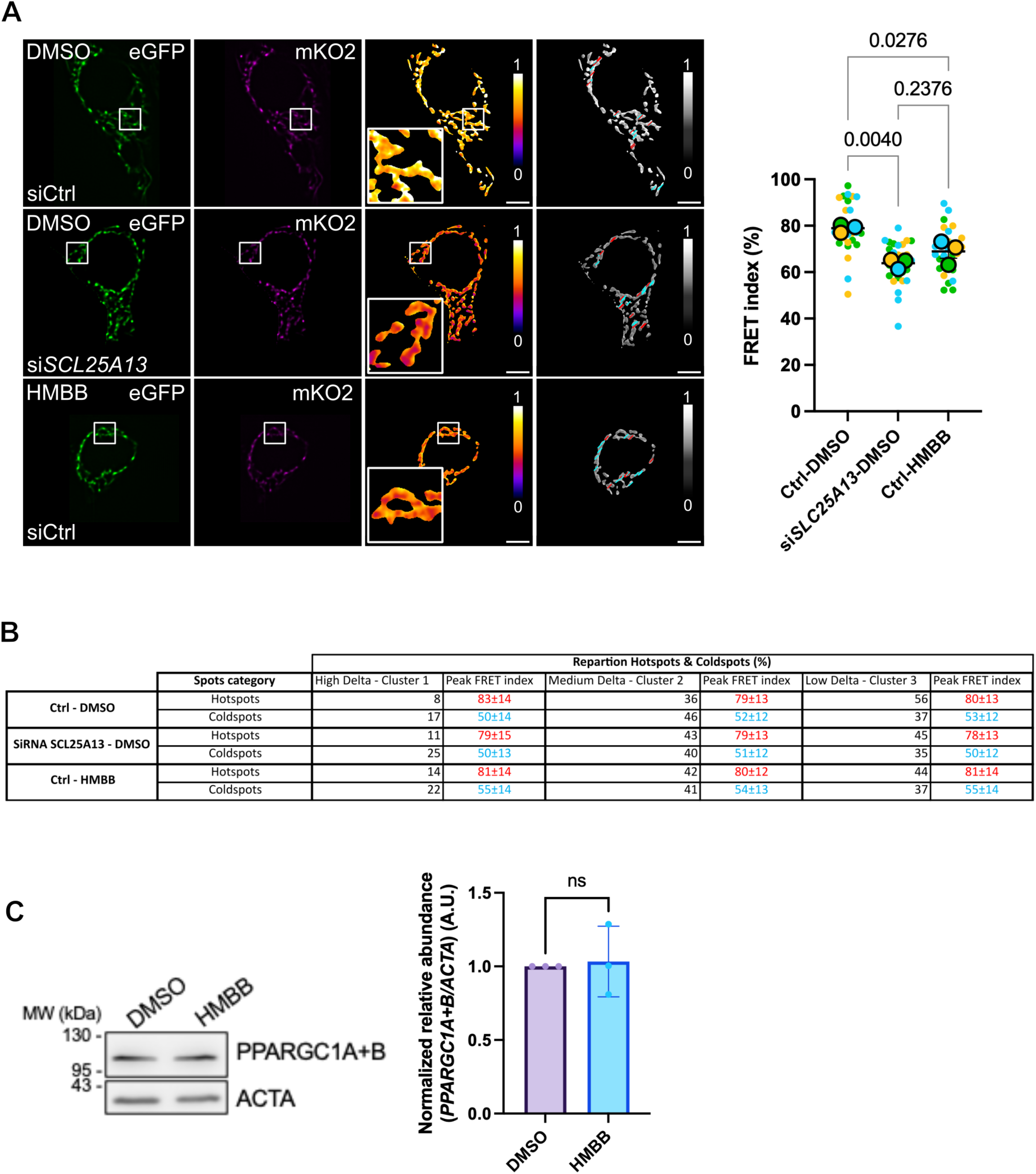
(Figure on the previous page) SLC25A13 abundance shapes mitochondrial metabolic heterogeneity. (**A**) (From left to right) eSRRF-reconstructed images of GFP, mKO2 (FRET), ratiometric FRET images and visual representation of ratiometric FRET shown in greyscale upon random line analyses of T47D cells expressing the mitoGO-ATeam2 biosensor and transfected with control or *SLC25A13*-specific siRNAs, and treated with DMSO or HMBB as indicated. Lines corresponding to FRET Hotspots or Coldspots are pseudocolored in red and cyan, respectively. Insets: higher magnification of the squared area. Graph: mean FRET index values (0-100%) in the indicated conditions. *n* = 10 cells per condition (small dots) in each of three biological replicates. Large dots indicate mean values for each biological replicate. Data are means ± S.D. Scale bar: 5 µm. (**B**) Table showing the overall repartition of FRET Coldspots and Hotspots detected with the MitoGO-ATeam2 biosensor in the indicated transfection condition, together with the quantity of FRET Coldspots and Hotspots depending on the degree of variation of the FRET index (clusters). For each cluster and spot type, the FRET index peak value is shown in red (FRET Hotspots) or cyan (FRET Coldspots). (**C**) Representative western blot and corresponding quantifications of the relative abundance of PPARGC1 in total lysates of T47D cells treated with DMSO or HMBB for 24 h. Loading control: ACTA. Data are from *n* = 3 independent experiments, and presented as means ± S.D. Individual values are presented as dots. A.U.: arbitrary units. \**P* < 0.05, \*\**P* < 0.01. ns: not significant.

BioSenSRRF line analyses revealed that the mitochondrial network in control cells was rather homogeneous in terms of ATP production. More than 50% of the high ATP-producing *loci* (hotspots) showed low FRET variability and belonged to cluster 3, whereas most of the low ATP-producing *loci* (coldspots) belonged to cluster 2 (Fig. 8B). Depletion of SLC25A13 dramatically increased the number of FRET hotspots and coldspots belonging to cluster 1. These FRET variations did not affect peak FRET values along the line, revealing that SLC25A13 knockdown reduced the size of ATP production *loci* and thereby induced ATP spatial heterogeneity without changing overall FRET levels (Fig. 8B). Among the selected AURKA/PHB2 partners, the effect on mitochondrial ATP patterning was unique to SLC25A13 depletion. Indeed, depletion of NDUFA9, ATP5F1A SAMM50 did not change hotspot or coldspot amounts, but reduced FRET index values in both cluster 2 and 3 hotspots and coldspots (Supplementary Fig. 8). This indicates a more homogeneous decrease in FRET distribution when ATP5F1A or SAMM50 are absent. Finally, we observed that the effect of *SLC25A13* siRNA was similar to that observed when treating cells with HMBB (Fig. 8A-B). This is due to the capacity of this compound to inhibit AURKA/PHB2-dependent mitochondrial turnover (Djehal *et al*., 2026) (Supplementary Fig. 6A), and not by lowering the levels of PPARGC1A+B (Fig. 8C).

Overall, AURKA and PHB2 regulate mitochondrial metabolism and mitochondrial ATP patterning through their proximity with a core IMM subset of proteins including SLC25A13, NDUFA9, and ATP5F1A.

## Discussion

AURKA is frequently overexpressed at the mRNA and protein levels in breast and hematological malignancies (Nikonova *et al*., 2013). AURKA has a wide variety of cellular functions and partners, and its overexpression changes cell physiology both at mitosis and during interphase (Nikonova *et al*., 2013; Bertolin & Tramier, 2020). The identification of AURKA within mitochondria further complicated this scenario, increasing the number of partners and pathways the kinase is involved in (Grant *et al*., 2018; Bertolin *et al*., 2018). Overexpressed AURKA can interact with PHB2 in mitochondria to trigger mitochondrial turnover by mitophagy (Bertolin *et al*., 2021). By activating mitophagy, a portion of the mitochondrial network is degraded, while the remaining part shows an increased metabolic activity. When inhibiting mitophagy using pharmacological compounds such as the natural polyphenol Xanthohumol (Bertolin *et al*., 2021), or the capsaicin analog HMBB (Djehal *et al*., 2026; Jolivet *et al*., 2026b), mitochondrial ATP production levels were restored to physiological levels. Despite this evidence, the molecular players linking mitophagy and mitochondrial metabolism remained unknown.

Here, we show that AURKA and PHB2 share a nexus of mitochondrial partners, including the aspartate/glutamate transporter SLC25A13, the import-related protein SAMM50, the Complex I subunit NDUFA9, and the Complex V subunit ATP5F1A. All these proteins play established roles in metabolism regulation. SLC25A13 helps maintain the NADH/NAD+ redox equilibrium between the cytosol and the mitochondrial matrix, both in physiology and in cancer cells (Rabinovich *et al*., 2020; Ruprecht & Kunji, 2020). SAMM50 controls the biogenesis and assembly of respiratory chain Complexes I, III and IV, by forming a supramolecular complex with MICOS subunits (Ott *et al*., 2012; Schaumkessel *et al*., 2025). Interestingly, mitochondrial Complex I subunits are strong interactors of AURKA and PHB2 (Fig. 1, 2, Supplementary Fig. 2, 3), while Complex IV subunits are upregulated by AURKA overexpression (Bertolin *et al*., 2021). Finally, NDUFA9 and ATP5F1A are key subunits of mitochondrial Complex I and V. The combination of MS/MS data with FRET analyses refined the identity of proteins within this metabolic network governed by AURKA and PHB2. In addition, it strongly suggests that AURKA controls mitochondrial metabolism directly, by acting through a metabolism-related set of proteins at the IMM. Among these proximal factors, it is interesting to note that none of the AURKA and PHB2 partners seem to be essential for AURKA/PHB2-dependent mitophagy regulation. *SLC25A13* knockdown was previously reported to activate the autophagy mediator LC3-II in melanoma cell lines (Rabinovich *et al*., 2020). However, this does not seem to be the case in T47D breast cancer cells. In this model, the abundance of PMPCB is similar to that in control cells, and mitophagy does not seem to be activated in the absence of SLC25A13 (Fig. 7B). Despite differences in organellar distribution in SLC25A13-depleted cells (Fig. 7A), no primary difference in mitochondrial mass was observed when NDUFA9, ATP5F1A, or SAMM50 were knocked down in the same model. This further supports the notion that mitophagy occurs upstream of metabolic reprogramming and relies only, among the protein-protein proximities explored, on the functional interaction between AURKA and PHB2. The nexus of proteins driving metabolic adaptation relies on a set of actors that are in spatial proximity to both AURKA and PHB2. This reinforces the tight coupling between mitophagy and metabolism in AURKA-overexpressing cells, and suggests that this spatial proximity may be advantageous for rapidly modulating metabolic rewiring upon mitophagy activation. Inhibition of mitophagy with HMBB treatment does not prevent proteins within this nexus from being in physical proximity, such as between AURKA or PHB2 and SLC25A13 (Supplementary Fig. 5B). As HMBB is a AURKA/PHB2-targeting compound with the capacity to catalytically inhibit the mitochondrial pool of AURKA (Djehal *et al*., 2026), our results suggest that AURKA kinase activity is not necessary to maintain metabolism-related protein vicinities. However, future studies will elucidate whether AURKA-dependent phosphorylation of specific proteins occurs within the nexus, or if AURKA and PHB2 act purely as scaffolds to drive metabolic adaptation.

HMBB also confirmed its strong relevance to restore mitochondrial metabolism (Jolivet *et al*., 2026b). At the proximal interactome level, treatment with HMBB shifts NDUFA9 and ATP5F1A proximal protein profiles back to those of control cells, despite AURKA overexpression (Fig. 6). This indicates that the effect of HMBB is not limited to individual proteins, but it extends over a significant portion of the NDUFA9 and ATP5F1A proximal interactomes. At the cellular level, HMBB restored metabolic heterogeneity, as measured by the mitoGO-ATeam2 biosensor in BioSenSRRF mode (Fig. 8). These results further corroborate the close relationship between mitophagy and metabolic rewiring upon AURKA overexpression. They also highlight the key role of AURKA in directly shaping metabolic heterogeneity. It is interesting to observe that T47D cells are metabolically homogeneous (Fig. 8A), while alternative breast cancer cells such as MCF7 are more heterogeneous (Jolivet *et al*., 2026b). In both cell types, AURKA directly maintains metabolic heterogeneity. This is achieved either through AURKA overexpression (Jolivet *et al*., 2026b), the inhibition of AURKA/PHB2-dependent mitophagy with HMBB, or the absence of key partners such as SLC25A13 (Fig. 8A). Both MCF7 and T47D cells rely on OXPHOS, with T47D cells showing the highest oxygen consumption rates (Sharma *et al*., 2023). Yet, it cannot be excluded that transient AURKA overexpression is an acute stress that leads to strong mitophagy activation, changes in mitochondrial ultrastructure (Fig. 3), and a short-term increase in mitochondrial heterogeneity as stress responses (Jolivet *et al*., 2026b). In contrast, T47D cells may have evolved to maintain not only high endogenous AURKA levels (Bertolin *et al*., 2018), but also to ensure constant turnover rates (Supplementary Fig. 6A), limiting overall ultrastructural and morphological changes (Fig. 5; Supplementary Fig. 6A-C), and maintaining a metabolically homogeneous network. We show that lowering the abundance of the nexus protein SLC25A13 is a direct way to re-establish metabolic heterogeneity by inducing primary metabolic dysfunction. Alternatively, this can be achieved indirectly through mitophagy inhibition after HMBB treatment, which might maintain metabolically dysfunctional mitochondria within the network.

In conclusion, our results identify SLC25A13, NDUFA9 and ATP5F1A as key proximal interactors of AURKA and PHB2 within the IMM. We show that these interactors drive metabolic adaptations in AURKA-overexpressing cells. Our findings also show that the AURKA/PHB2-targeting compound HMBB restores AURKA-dependent metabolic rewiring and spatial metabolic heterogeneity. This paves the way for the use of HMBB to induce metabolic heterogeneity, or the development of compounds targeting SLC25A13, NDUFA9, or ATP5F1A to counteract the metabolic consequences of AURKA overexpression in cancer.

## Limitations of the study

While our study identifies a platform of inner mitochondrial membrane proteins that couple AURKA and PHB2 to the control of ATP production and organelle architecture, several limitations should be acknowledged. First, our proximity interactomics and live-cell imaging were primarily conducted in HEK293, MCF7, and T47D cell lines, which may not fully recapitulate the complexity of mitochondrial regulation in all cancer types or *in vivo* contexts. Second, the use of FRET/FLIM microscopy and proximity labeling captures stable or transient interactors within a < 10 nm (FFET/FLIM) or 10-20 nm radius (BioID/TurboID), potentially missing more distant or weakly associated partners. Third, although we demonstrate that AURKA overexpression rewires the interactomes of NDUFA9 and ATP5F1A, the functional consequences of these rewiring events on individual respiratory complex activities were not directly measured here. While HMBB restores interactomes and ATP heterogeneity, its precise mechanism of action on downstream metabolic pathways requires further investigation, including whether its modulation of AURKA kinase activity is essential for maintaining protein-protein proximities. Finally, the role of SLC25A13 in sustaining metabolic heterogeneity was inferred from knockdown experiments, and future studies should explore its contribution in more physiological or pathological settings, such as in animal models or patient-derived samples.

## Materials and methods

### Expression vectors and molecular cloning

The PHB2 CDS was cloned into pcDNA5 FRT/TO C-ter BirA*Flag vectors for BioID, while ATP5F1A and NDUFA9 CDS were cloned into a pcDNA5 FRT/TO C-ter TurboID-3xFlag using the AscI/NotI restriction sites. The COX8 mitochondrial targeting signal (MTS) was cloned into a TurboID-3xFlag and used as a control. The full list of plasmids used in the study is in Supplementary Table 1.

### Cell culture procedures, chemicals

MCF7 (HTB-22) and T47D (HTB-133) cells were purchased from the American Type Culture Collection, while Flp-In™ T-REx™ HEK293 cells were purchased from Thermo Fisher Scientific. All cell lines were kept mycoplasma-free throughout experiments and checked monthly for potential mycoplasma contamination. Cells were grown in Dulbecco’s Modified Eagle’s Medium (DMEM) already containing 1% L-glutamine (Thermo Fisher Scientific), and supplemented with 10% FBS (Biosera) and 1% penicillin-streptomycin (Thermo Fisher Scientific). For all live microscopy experiments, cells were grown at 37 °C either in Nunc Lab-Tek Chamber slides (Thermo Fisher Scientific) or in polymer-coated µ-Slide 8 Well high slides (Ibidi). Before FLIM imaging, standard growth media was replaced with phenol red-free Leibovitz’s L-15 medium (Thermo Fisher Scientific) supplemented with 20% FBS and 1% penicillin–streptomycin. For stable transfections, Flp-In™ T-REx™ HEK293 cells were co-transfected with BirA* or TurboID tag-encoding constructs and pOG44, and then selected with 200 µg/ml Hygromycin B (Thermo Fisher Scientific). Transient plasmid transfections were performed with Lipofectamine 2000 (Thermo Fisher Scientific), while siRNA transfections were performed with Lipofectamine RNAiMax according to the manufacturer’s instructions. Transient transfection mixes were incubated for 48 h before microscopy analyses, cell fractionation experiments, or total cell extraction for western blotting analyses. Allstars negative control (SI03650318) siRNAs, a functionally-verified siRNA against *SAMM50* (SI03019079), a siRNA against *SLC25A13* (SI05012518), *ATP5F1B* (SI02626722), *ATP5F1A* (SI04989873), and *NDUFA9* (SI05023088) were purchased from Qiagen. HMBB was resuspended in DMSO and stored at −20 °C for long-term storage. It was used at a final concentration of 50 µM, and incubated for 6 h before mitochondrial fractionation for TurboID experiments or for 24 h before live cell imaging or protein extraction experiments.

### Immunocytochemistry, confocal and dSTORM super-resolution microscopy

T47D cells were fixed in 4% paraformaldehyde (Euromedex), stained using standard immunocytochemical procedures, and mounted in ProLong Gold Antifade reagent (ThermoFisher Scientific). Mitochondria were stained with a polyclonal rabbit anti-PMPCB (16064–1-AP; Proteintech) used at a 1:500 dilution and a secondary anti-rabbit antibody coupled to Alexa Fluor488 used at a 1:2000 dilution (ThermoFisher Scientific). Images were acquired with a Leica SP8 inverted confocal microscope (Leica) driven by the Leica Acquisition Suite (LAS) software and a 63x oil immersion objective (NA 1.4).

For dSTORM microscopy, MCF7 and T47D cells were fixed in 4% paraformaldehyde (Euromedex) and 0.2% glutaraldehyde, stained using standard immunocytochemical procedures, and mounted with a home-made buffer previously described in (Beghin *et al*., 2017) and composed of 120 μg/ml Pyranose Oxidase (Sigma-Aldrich, P4234) and 57 μg/ml catalase (Sigma-Aldrich, C3515). β-Mercaptoethylamine (= cysteamine, Sigma-Aldrich 30070) was prepared as a 1 M stock solution in water and stored at 4°C. Tris HCl (20 mM Tris, 50 mM NaCl) pH 7.2 was used as a base for the buffer. Primary antibodies for dSTORM were: mouse ATP5F1B (Santa Cruz, sc-166462), used at 1:500, and Aurora A (clone 5C3 (Cremet *et al*., 2003)) used at a 1:50 final dilution; rabbit PHB2/REA (ThermoFisher Scientific, PA5-79817) used at 1:500, and MIC60 (Abcam, ab137057) at 1:500. Secondary antibodies were Alexa Fluor647 (Thermo Fisher Scientific) used at 1:5000, and CF 532 (Biotium) used at 1:2000. Acquisitions were made with an Abbelight super resolution microscope (Abbelight) constituted by an Olympus IX83 microscope, an Abbelight SAFe 180 nanoscopy module, a focus hold ZDC2 system, two lasers at 640 and at 532 nm (Oxxius), a pco.panda sCMOS camera (PCO AG), equipped with a 100X oil-immersion objective (NA 1.5), and driven by the NEO software (Abbelight, France).

### FLIM and ratiometric FRET imaging

In FRET/FLIM experiments, GFP was used as a FRET donor in all experiments, and its decrease was measured as in (Djehal *et al*., 2026). FLIM analyses were performed with a time-gated custom-built system coupled to a Leica DMI6000 microscope (Leica) and equipped with a CSU-X1 spinning disk module (Yokogawa) and a 63x oil immersion objective (NA 1.4), a picosecond pulsed supercontinuum white laser at 40 MHz frequency (Fianium), and a High-Rate Intensifier (LaVision) coupled to a CoolSNAP HQ2 camera (Roper Scientific). GFP was excited at 480±10 nm, and fluorescence emission was selected with a band-pass filter (483/35 nm) (Semrock). To calculate fluorescence lifetime, five temporal gates with a step of 2 ns each allowed the sequential acquisition of five images covering a total delay time from 0 to 10 ns. The FLIM setup was controlled by the Inscoper Suite solution (Inscoper), which also allowed for lifetime measurements and calculations in real-time during acquisition. Lifetime was calculated only when pixel-by-pixel fluorescence intensity in the first gate was above 2,000 grey levels. Raw lifetime values were extracted by analyzing images post-acquisition with the Fiji software (NIH). To calculate ΔLifetime values, the mean lifetime of the cells in the donor-only condition (AURKA-GFP + Empty vector or PHB2-GFP + Empty vector) was calculated and then used to normalize data in all the analyzed conditions and for each independent experiment, as previously reported (Bertolin *et al*., 2020).

For ratiometric FRET imaging, images were acquired using an inverted DMi8 microscope (Leica) equipped with a spinning disk CSU-X1 (Yokogawa) module, a Photometrics Evolve EmCCD camera, a 63x (NA 1.4) oil immersion objective, and controlled by the Inscoper Suite solution. The excitation and emission wavelengths for cpGFP were 488 and 525/40 nm; a 488 nm excitation and a 607/36 nm emission were used for mKO2. For each channel, 200 frames were acquired with an exposure time of 25 ms to image the donor and the acceptor of MitoGO-ATeam2. Image processing was described in (Jolivet *et al*., 2026b). Briefly, raw images were processed using the Fiji software (NIH) with a publicly available custom-made macro (https://github.com/cyclochondria/BioSenSRRF.git). The macro is divided into multiple steps to facilitate image treatment. First, each raw image undergoes background subtraction. To this end, images are thresholded to delete pixels with intensity below 500 grey levels. Images are then reconstructed with the eSRRF plugin (Laine *et al*., 2023) using settings predetermined with the *parameter sweep* function. In our case, the following settings were used: *Magnification=5, Radius=3, Sensitivity=3, Number of frames=200, Average reconstruction*. The mitochondrial area was determined using the auto-threshold function in Li mode on the cpGFP channel. The FRET index after eSRRF reconstruction was calculated by dividing the fluorescence intensity on the acceptor channel (mKO2) by the intensity of the donor channel (GFP) in the GFP-positive area. FRET images were produced using the *Image Calculator* tool embedded within Fiji.

The random line analysis was performed on FRET images after SRRF reconstruction using Fiji (https://github.com/cyclochondria/BioSenSRRF.git). Each line was set at a dimension of 25 pixels and is inclined at 45°. For each image, 50 random lines were generated, and the FRET index on each line was then calculated. The lines were filtered and classified in clusters following the same procedure used in (Jolivet *et al*., 2026b). The lines showing a Gaussian and an Inverse Gaussian profile were extracted according to their curvature calculated through their discrete second derivative, and the percentage of distribution of Gaussian (FRET Hotspots) and Inverse Gaussian (FRET Coldspots) were plotted for each replicate. The distribution of FRET Hotspots and Coldspots was then classified according to the variation of FRET along the line (High, medium, and low variations) for each replicate. The random line generation and filtration pipeline was generated on RStudio (https://github.com/cyclochondria/BioSenSRRF.git) (Jolivet *et al*., 2026b).

### Mitochondrial morphology and relative area analyses

The excitation/emission wavelengths for Alexa Fluor488 were 488 and 525/50 nm, respectively. Relative mitochondrial area (A_PMPCB_) calculations were obtained with the Fiji software (NIH) on maximal projections of confocal images acquired as above. As previously reported (Bertolin *et al*., 2021), A_PMPCB_ was determined as the ratio between the area covered by PMPCB selected with an automatic threshold mask and the total cell area. Mitochondrial length and branching were estimated with aspect ratio and form factor calculations, extracted from confocal images acquired as above. Calculations were performed as in (Koopman, 2005).

### Western blotting

Total protein fractions were obtained by lysing cells in 50 mM Tris-HCl (pH 7.5), 150 mM NaCl, 1.5 mM MgCl_2_, 1% Triton X-100, and 0.5 mM dithiothreitol (DTT) supplemented with 0.2 mM Na_3_VO_4_, 4 mg/ml NaF, 5.4 mg/ml b-glycerophosphate and protease inhibitors (Complete Cocktail, Roche) followed by centrifugation at 13,000 g for 20 min at 4°C. All protein fractions were assayed using the Bradford reagent (Bio-Rad) and then boiled in Laemmli sample buffer, resolved by SDS–PAGE, transferred onto a nitrocellulose membrane (GE Healthcare), and analysed by western blotting. Primary antibodies were as follows: mouse monoclonal anti-Actin (Abcam, 8224) used at 1:250, rabbit polyclonal anti-PMPCB (16064–1-AP; Proteintech) used at 1:1000, anti-SLC25A13 (Abcam, ab96303) used at 1:1000, rabbit monoclonal anti-PPARGC1A+B (Abcam, ab188102) used at 1:1000, rat anti-Tubulin alpha clone YL1/2 (MAB1864 Millipore) used at 1:5000. Secondary horseradish-peroxidase-conjugated antibodies (anti-mouse and anti-rabbit) were purchased from Jackson ImmunoResearch Laboratories; anti-rat antibodies were purchased from Bethyl Laboratories. The membranes were incubated with commercially available (Pierce) enhanced chemiluminescence substrate. Chemiluminescence signals were captured on an Amersham Imager 680 (GE Healthcare). Integrated band densities were quantified with Fiji (NIH). The relative abundance of specific bands of interest was calculated by normalizing them towards the abundance of respective loading controls and indicated in each graph.

### BioID and TurboID experiments

For BioID, three separate replicates of 10 cm plates containing cells at 60% confluence were incubated for 24 hours in complete media with 1 μg/ml tetracycline (Sigma) and 50 μM biotin (Thermo Fisher Scientific). Cells were harvested, pelleted at 300 ×g for 3 minutes, washed twice with PBS, and the dried pellets were quickly frozen.

For TurboID, three separate replicates of cells transfected with pcDNA3.1 or pcDNA3 AURKA-6xHis were performed in 10 cm plates containing cells at 80% confluence. Cells were incubated for 24 hours in complete media with 1µg/ml tetracycline (Sigma-Aldrich) and 6 hours with DMSO or HMBB at 50 µM, and biotin at 50 µM (Thermo Fisher Scientific). The cells were then washed and harvested in 1X PBS (Euromedex), then pelleted at 300 ×g for 3 minutes. We then performed mitochondrial fractionation as previously described (Bertolin *et al*., 2018). Briefly, cells were lysed in 210mM mannitol, 70mM sucrose, 5mM Tris pH 7.4, 0.2 mM EGTA, 0.1mM EDTA, 0.1% BSA, 0.2 mM Na_3_VO_4_, 0.5mM DTT, 4mg/ml NaF, 5.4mg/ml Beta-GlycerolPhosphate, 1X protease inhibitors (Roche) followed by centrifugation at 13,000 g for 30 min at 4°C.

Pellets were resuspended in 10mM HEPES KOH pH 7.4, 250mM Sucrose, 0.5mM EGTA, 2mM EDTA, 1mM DTT, and frozen at -80°C before dosing.

For TurboID and BioID, pellets containing mitochondrial and total fractions, respectively, were then resuspended in lysis buffer containing 50 mM Tris-HCl pH 7.5, 150 mM NaCl, 1 mM EDTA, 1 mM EGTA, 1% Triton X-100, 0.1% SDS, 1:500 protease inhibitor cocktail (Sigma-Aldrich), 1:1000 Turbonuclease (BPS Bioscience), and rotated end-over-end at 4 °C for 1 hour. The mixture was briefly sonicated to break up visible aggregates and then centrifuged at 45,000 × g for 30 minutes at 4 °C. 25 μl of packed, pre-equilibrated Streptavidin Ultralink Resin (Pierce) was added, and the mixture was rotated for 3 hours at 4 °C. The beads were pelleted by centrifugation at 300 × g for 2 minutes and transferred to a new tube with 1 mL of lysis buffer. The beads were washed once with lysis buffer, and four times with 50 mM ammonium bicarbonate (pH 8.3). Tryptic digestion was carried out by incubating the beads with 1 μg MS-grade TPCK trypsin (Promega, Madison, WI) dissolved in 200 μl of 50 mM ammonium bicarbonate (pH 8.3) overnight at 37 °C. The following day, an additional 0.5 μg MS-grade TPCK trypsin was added to the beads and incubated for another 2 hours at 37 °C. After centrifugation at 2000 × g for 2 minutes, the supernatant was collected and transferred to a new tube. Two more washes were performed with 50 mM ammonium bicarbonate and combined with the initial eluate. The sample was freeze-dried and resuspended in buffer A (2% ACN, 0.1% formic acid). One-third of each BioID sample and 1/8^th^ of each TurboID sample was analyzed per each mass spectrometer run.

In PHB2-BioID experiments, protein identification was achieved by comparing all MS/MS data with the *Homo sapiens* proteome database (Uniprot, March 2020 release, Canonical+Isoforms, including 42,360 entries plus manually added viral bait protein sequences), utilizing MaxQuant software version 1.5.8.3. Trypsin was used for digestion, allowing up to 2 missed cleavages. Methionine oxidation and N-terminal protein acetylation were set as variable modifications. Label-free quantification (LFQ) was conducted using the software’s default settings with an initial mass tolerance of 6 ppm in MS mode and 20 ppm for MS/MS fragmentation data. An initial mass tolerance of 6 ppm was chosen for MS mode, while 20 ppm was used for MS/MS fragmentation data. Protein and peptide identification parameters were set with a 1% false discovery rate (FDR) and required at least 2 unique peptides per protein. For control runs (FlagBirA* alone samples), the LFQ values were reduced to the top three for each ID, forming the control group for comparison with PHB2-BirA*Flag triplicates. The LFQ data underwent log2(x) transformation.

For TurboID samples, raw data were processed using DIA-NN software (version 1.9.2). A library-free approach was utilized to search the UniProt reviewed *Homo sapiens* database (September 2024, 20,420 entries). The search parameters allowed up to two missed cleavages during trypsin digestion, with methionine oxidation designated as a variable modification. Peptides were filtered based on a length range of 7–30 amino acids, a precursor charge range of 2–4, a precursor m/z range of 300–1300, and a fragment ion m/z range of 100–1700. False discovery rates (FDRs) were maintained at 1% for both protein and peptide levels. The “match between runs” feature was activated, and the quantification strategy was set to “robust LC (high accuracy)”.

For group comparisons, abundance comparisons were performed between the three biological replicates of each group using a Student’s *t*-test, requiring at least three out of three experimental values in one of the two groups. LFQ values were log2-transformed before statistical analysis. When no values were detected in one group, missing values were replaced by the minimal detected value of each run to avoid division by zero. When only one value was detected in a group, two synthetic values were added at +1 and −1 log2 LFQ around the detected value to allow stringent filtering. When two values were detected, the third value was imputed as the average of the two experimental values. No additional normalization or batch correction was applied to the raw analysis.

For the PHB2 BioID experiment, only preys detected in all three PHB2-BirAFlag bait replicates were retained for downstream analysis. Control LFQ values from FlagBirA alone samples were reduced to the top three values for each protein ID to define the control group used for comparison with the PHB2-BirA*Flag triplicates. High-confidence proximal interactors were defined using a p-value cut-off of 0.01 and a log2 fold-change > 1 versus control. All experimental values are provided in Supplementary Table 3.

For TurboID experiments, high-confidence proximal interactors were similarly identified using a p-value cut-off of 0.01 and a log2 fold-change > 1 versus control. All experimental values are provided in Supplementary Tables 4-7.

### Statistical analyses

Two-way ANOVA with Tukey’s multiple comparison test was used to compare the effect of the transfection and of the HMBB treatment on ΔLifetime efficiencies (Supplementary Fig. 5B), the relative mitochondrial area (Supplementary Fig. 6A), and on mitochondrial morphology (Supplementary Fig. 6B-C). Ordinary one-way ANOVA with Šídák’s multiple comparison test was used to compare the effect of the transfection on the relative mitochondrial area (Fig. 7A) and on the FRET index (Fig. 8A, Supplementary Fig. 8A). Unpaired *T*-tests with Welch’s correction were used to compare the effect of the transfection condition (Fig. 2, Supplementary Fig. 2 and 3), on colocalization efficiencies (Fig. 3 and 5), and on protein abundance (Fig. 7B-C, Fig. 8C). Statistics were performed under GraphPad Prism v. 10. Statistics for biological replicates were performed using SuperPlotsofData under GraphPad Prism v. 10 (Lord *et al*., 2020).

## Author Contributions

CRediT taxonomy: Conceptualization (G.B.), Data Curation (C.C., N.J., G.B.), Formal Analysis (C.C., N.J., D. K., G.B.), Funding Acquisition (G.B.), Investigation (C.C., N.J., D. K., G.B.), Methodology (C.C., N.J., D. K., E.C., G.B.), Project Administration (G.B.), Resources (E.C., G.B.), Software (N.J.), Supervision (G.B.), Validation (G.B.), Visualization (C.C., N.J., G.B.), Writing – original (G.B.), Writing – review and editing (C.C., N.J., D. K., E.C., G.B.)

## Acknowledgments

The authors wish to thank Malvina Salami for technical assistance. HMBB was kindly provided by Laurent Désaubry (Univ. Strasbourg, France). We would also wish to thank Stéphanie Dutertre and Xavier Pinson at the Microscopy Rennes Imaging Center (MRic, BIOSIT, Biogenouest) for assistance with fluorescence microscopy experiments, and Arthur Masson (INRIA, Rennes, France) for help with the GCoPS software. MRic is a member of the national infrastructure France-BioImaging, supported by the French National Research Agency (ANR-24-INBS-0005 FBI BIOGEN). We would like to thank the OrganOmics Core Facility of the University of Lille, which is accredited and funded by the GIS IBiSA and is a member of the French Proteomics Infrastructure (ProFI) UAR 2048. We thank all the members of our team for constructive discussions and helpful comments.

## Data availability

Source microscopy data are available on Zenodo (https://zenodo.org/records/21333343 and https://zenodo.org/records/21342346) (Caron *et al*., 2026; Caron & Bertolin, 2026). Source BioID and TurboID data will be made publicly available upon publication. Fiji/ImageJ and RStudio custom-made macros for BioSenSRRF analyses are available on https://github.com/cyclochondria/BioSenSRRF.git. Standalone versions of the BioSenSRRF software are available on Zenodo (Jolivet *et al*., 2026a) (https://zenodo.org/records/18266611)

All other data are available from the corresponding author (G.B.) upon request.

