## Supplementary Material v2 for "Mitochondrial inner membrane partners of Aurora kinase A/AURKA and PHB2 shape organelle metabolic heterogeneity"

This file contains Supplementary Figures 1-9.

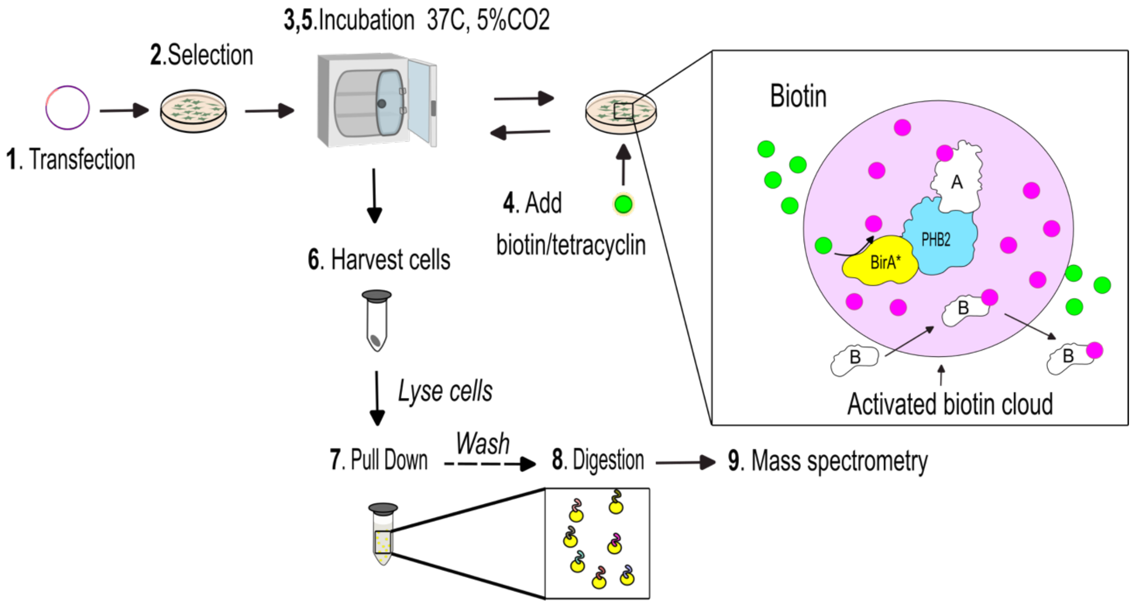


**Supplementary Fig. 1. Experimental pipeline of PHB2 BioID*.** (**1**) Flp-In™ T-REx™ HEK293 cells are transfected with pcDNA5 FRT/TO C-ter BirA*Flag vectors. (**2**) Stable clones are selected Hygromycin B and maintained (**3; 5**) at 37°C and in a 5% $CO_2$ atmosphere. (**4**) Tetracycline is added to induce the expression of PHB2-BirA*, while biotin is added to label stable (A) and transient (B) interactors. Cells are then harvested (**6**) and subjected to pull-down (**7**) before peptide digestion (**8**) and Mass Spectrometry analyses (**9**).

AURKA-GFP as donor

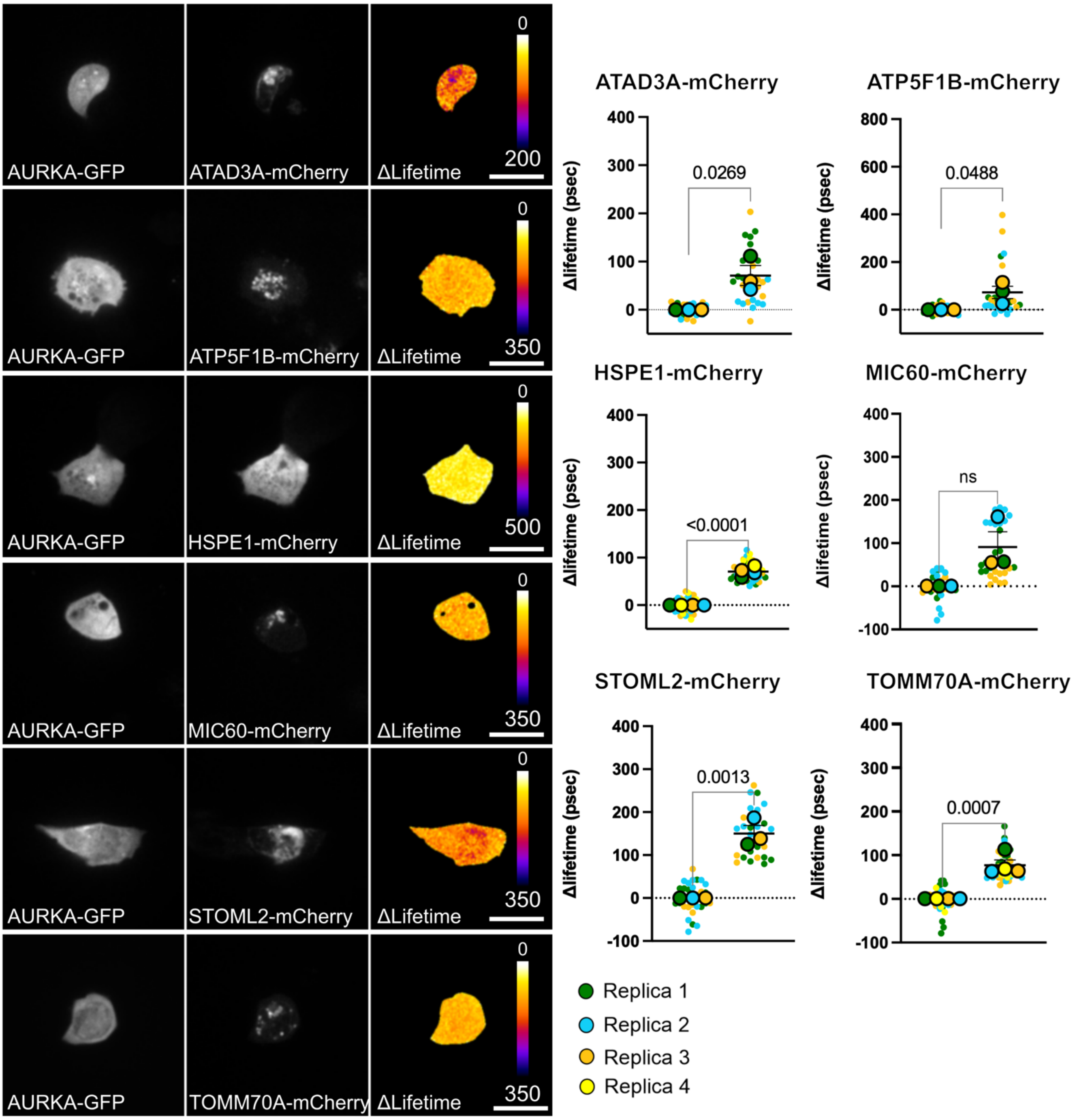


**Supplementary Fig. 2. AURKA shows proximity with IMM proteins.** Fluorescence, FRET/FLIM micrographs (left) and corresponding quantitative analyses (right) of MCF7 cells co-expressing AURKA-GFP (donor), and each of the indicated IMM proteins fused to mCherry (acceptors). Pseudocolor scale: pixel-by-pixel ΔLifetime. FRET measurements were performed in the acceptor-rich area. Scale bars: 5 µm. *n* = 10 cells per condition (small symbols) in each of three or four biological replicates. Large symbols indicate mean values for each biological replicate. Data are means ± S.D. Exact *P*-values are indicated for each comparison. ns: not significant.

PHB2-GFP as donor

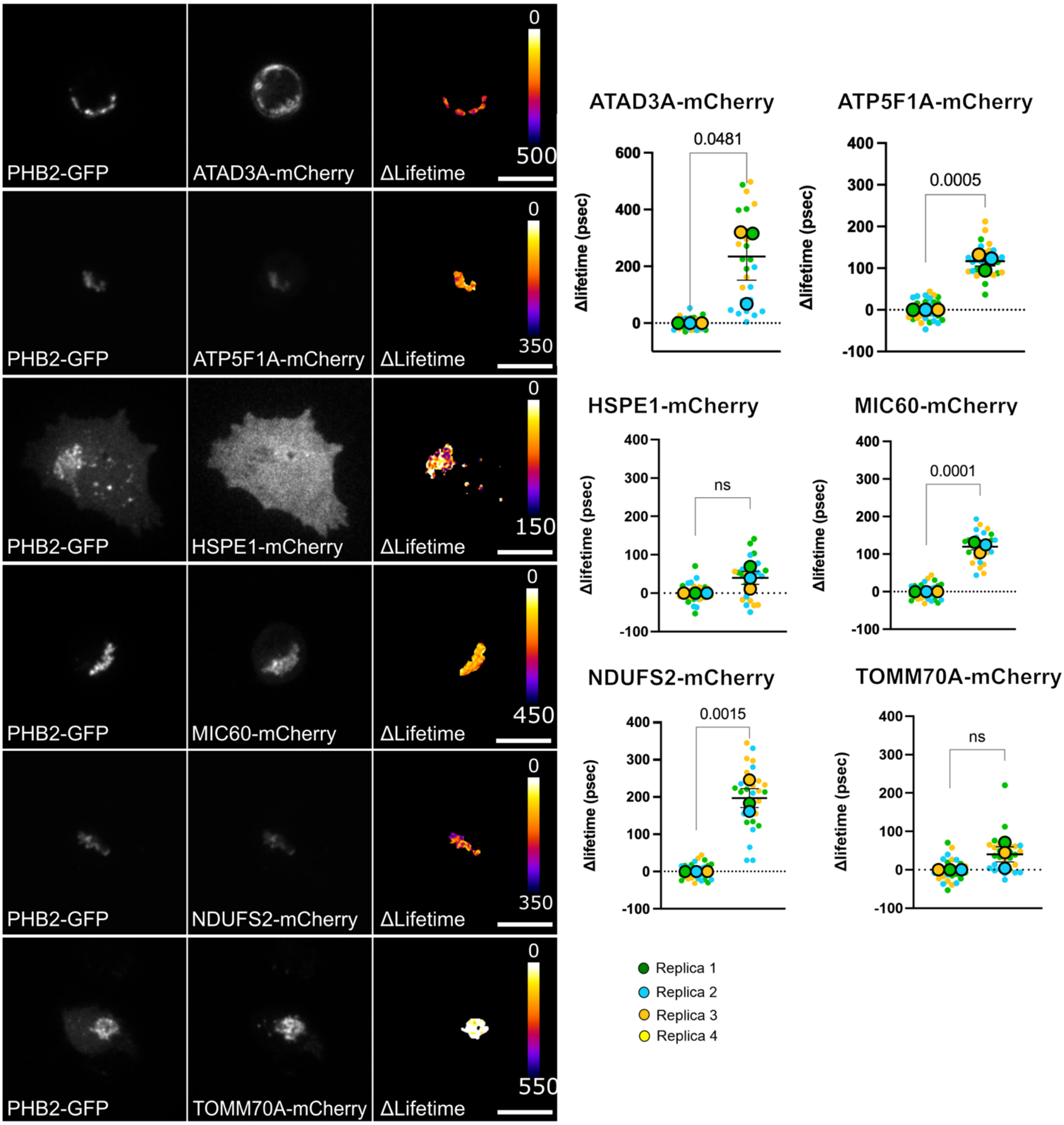


**Supplementary Fig. 3. PHB2 shows proximity with IMM proteins.** Fluorescence, FRET/FLIM micrographs (left) and corresponding quantitative analyses (right) of MCF7 cells co-expressing PHB2-GFP (donor), and each of the indicated IMM proteins fused to mCherry (acceptors). Pseudocolor scale: pixel-by-pixel ΔLifetime. FRET measurements were performed in the acceptor-rich area. Scale bars: 5 µm. *n* = 10 cells per condition (small symbols) in each of three or four biological replicates. Large symbols indicate mean values for each biological replicate. Data are means ± S.D. Exact *P*-values are indicated for each comparison. ns: not significant.

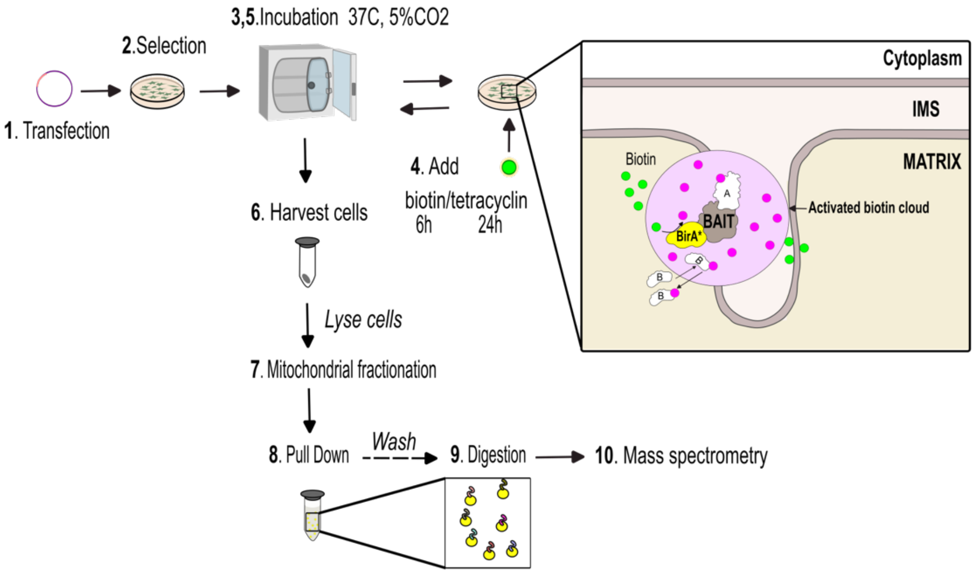


**Supplementary Fig. 4. Experimental pipeline for NDUFA9 and the ATP5F1A TurboID.** (**1**) Flp-In™ T-REx™ HEK293 cells are transfected with a pcDNA5 FRT/TO C-ter TurboID-3xFlag vectors. (**2**) Stable clones are selected Hygromycin B and maintained (**3; 5**) at 37°C and in a 5% $CO_2$ atmosphere. (**4**) Tetracycline is added to induce the expression of NDUFA9- or ATP5F1A-BirA*, while biotin is added to label stable (A) and transient (B) interactors. Cells are then harvested (**6**) and subjected to mitochondrial fractionation (**7**) followed by pull-down (**8**), peptide digestion (**9**) and Mass Spectrometry analyses (**10**).

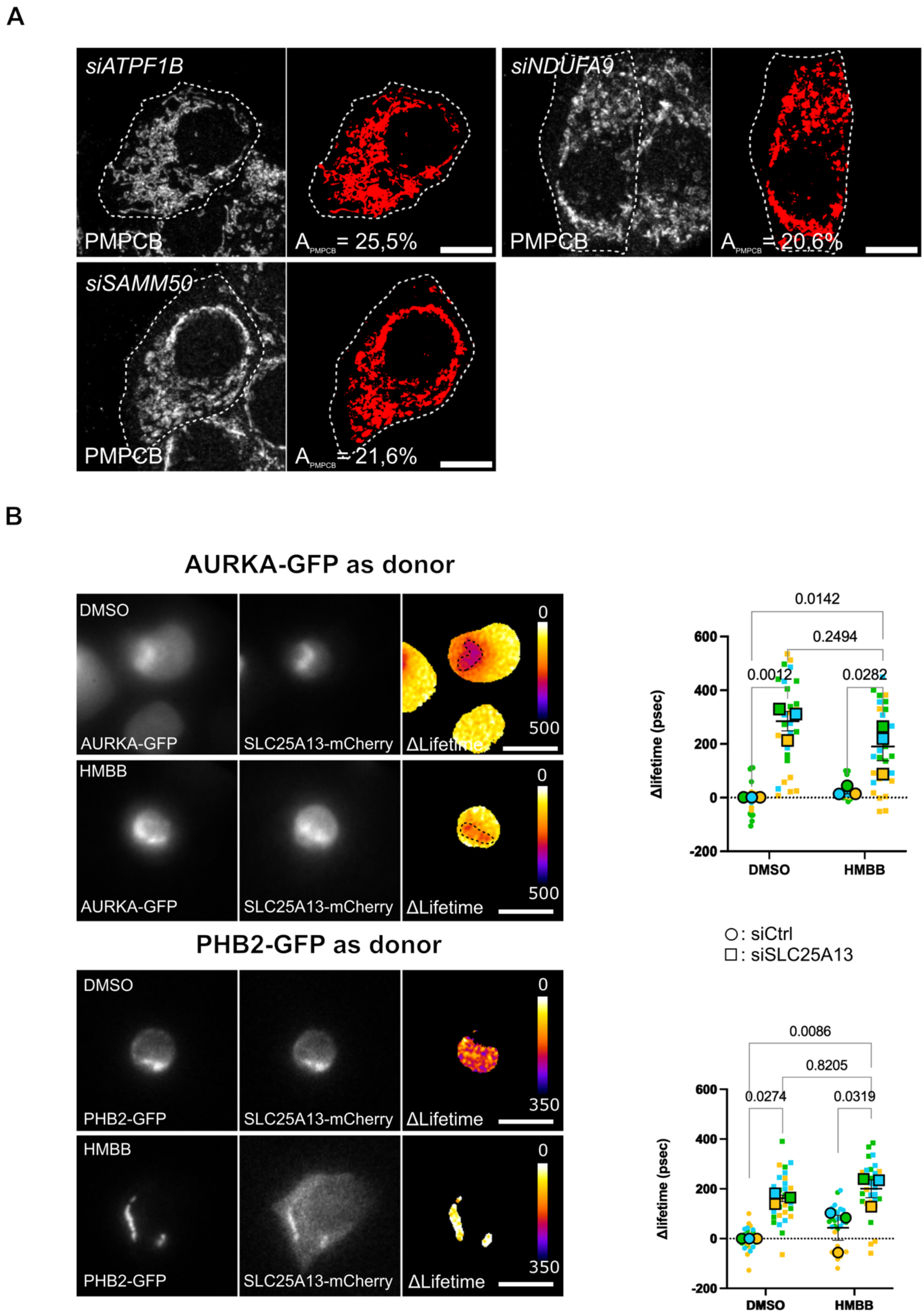


**Supplementary Fig. 5. Depletion of *ATP5F1A*, *SAMM50* and *NDUFA9* does not alter the relative mitochondrial area, while HMBB does not alter protein proximities between AURKA, PHB2, and SLC25A13.** (**A**) Relative mitochondrial area shown as the amount of PMPCB staining (threshold mask and corresponding quantification) in T47D cells transfected as indicated. $A_{PMPCB}$: mitochondrial area normalized against total cell area

(%). Scale bars: 10 µm. (**B**) Fluorescence, FRET/FLIM micrographs (left) and corresponding quantitative analyses (right) of MCF7 cells co-expressing either AURKA-GFP or PHB2-GFP (donors), and SLC25A13-mCherry (acceptor) and treated with DMSO or HMBB for 24 h. Pseudocolor scale: pixel-by-pixel ΔLifetime. FRET measurements were performed in the acceptor-rich area. Scale bars: 5 µm. $n$ = 10 cells per condition (small symbols) in each of three biological replicates. Large symbols indicate mean values for each biological replicate. Data are means ± S.D. Exact $P$-values are indicated for each comparison.

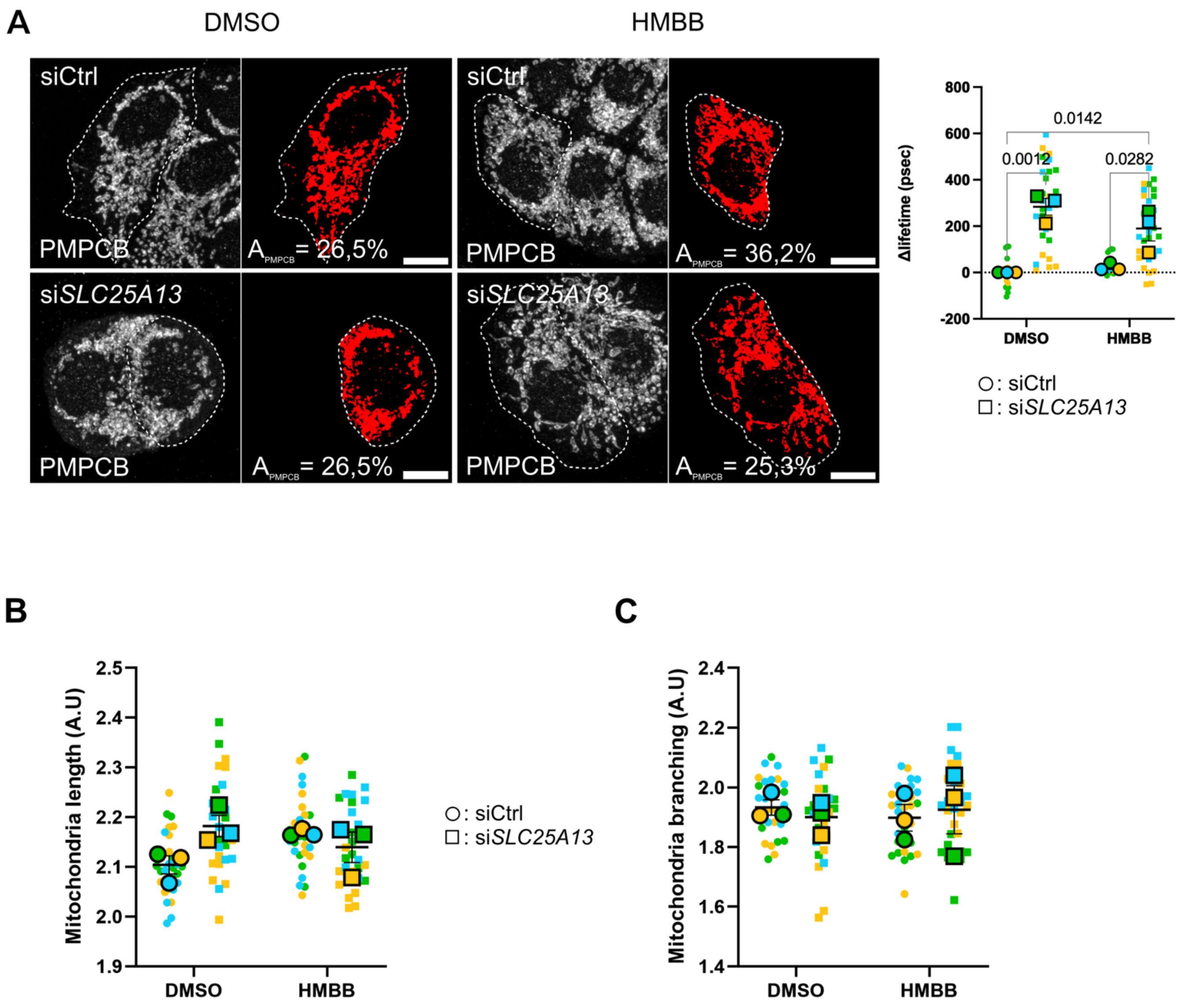


**Supplementary Fig. 6. HMBB restores mitochondrial mass and distribution, but does not alter mitochondrial length and branching.** (**A**) Relative mitochondrial area shown as the amount of PMPCB staining (threshold mask and corresponding quantification), and (**B**) mitochondrial length and (**C**) branching analyses performed in T47D cells transfected as indicated and treated with DMSO or HMBB for 24 h. $A_{PMPCB}$: mitochondrial area normalized against total cell area (%). Scale bars: 10 µm. *n* = 10 cells per condition (small dots) in each of three biological replicates. Large symbols indicate mean values for each biological replicate. Data are means ± S.D. Exact *P*-values are indicated for each comparison. All other comparisons were not significant.

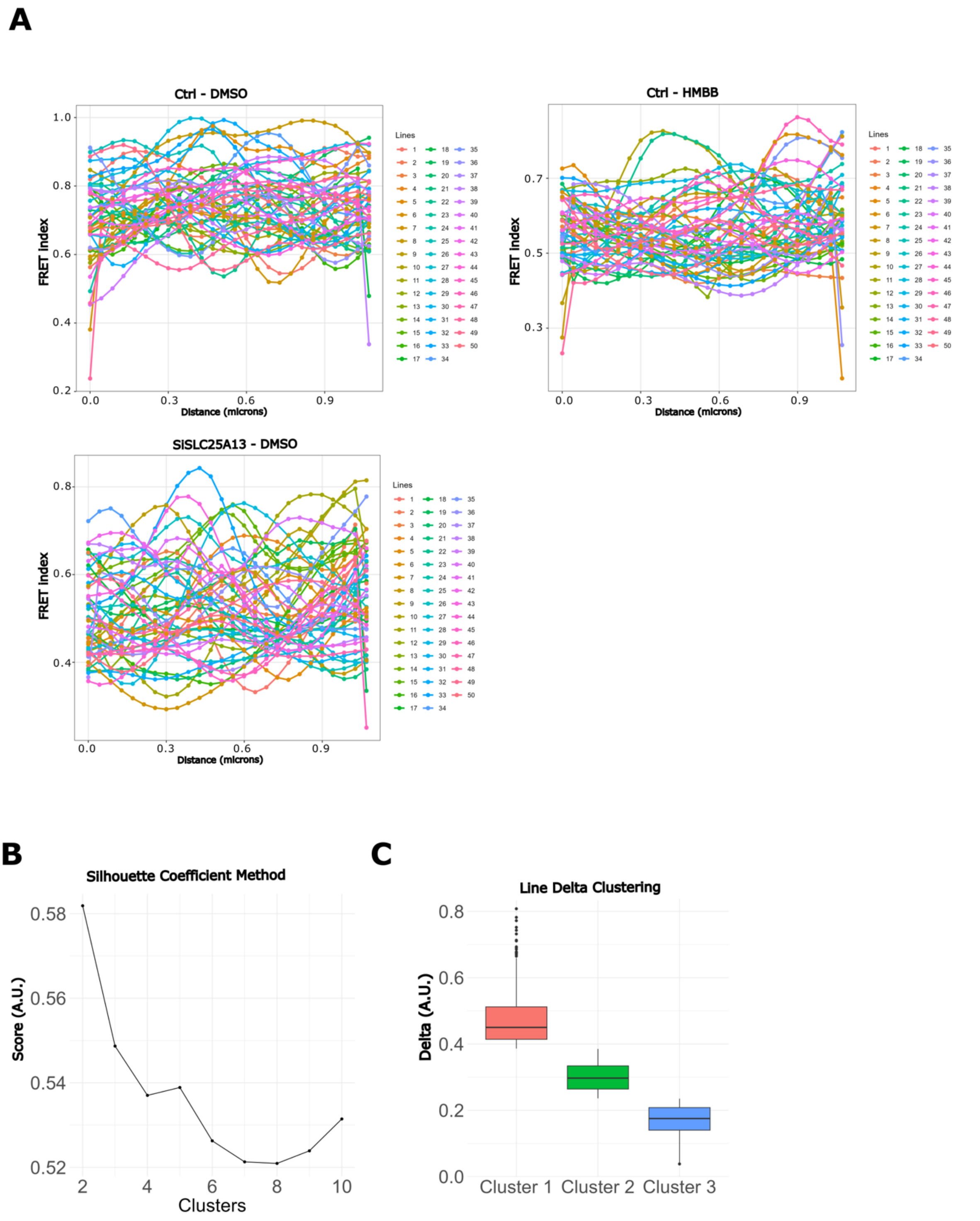


**Supplementary Fig. 7. Random line analyses and clusters of MitoGO-ATeam2 for BioSenSRRF analyses.** (**A**) Representative individual lines issued from the random line analyses in T47D cells transfected as indicated and treated with DMSO or HMBB for 24 h. The graphs show FRET index values (0-1) over distance (in µm) for each indicated condition. Each color represents an individual random line. (**B**) Mean silhouette coefficient score (A.U.) as a function of the number of clusters. (**C**) Distribution of the delta of FRET index values across the three identified clusters. Data go from mean to max, and boxplots show the 1$^{st}$ and 3$^{rd}$ interquartile range and the median delta value. Clusters are ordered by decreasing median delta values, highlighting differences in intensity variation among groups.

A

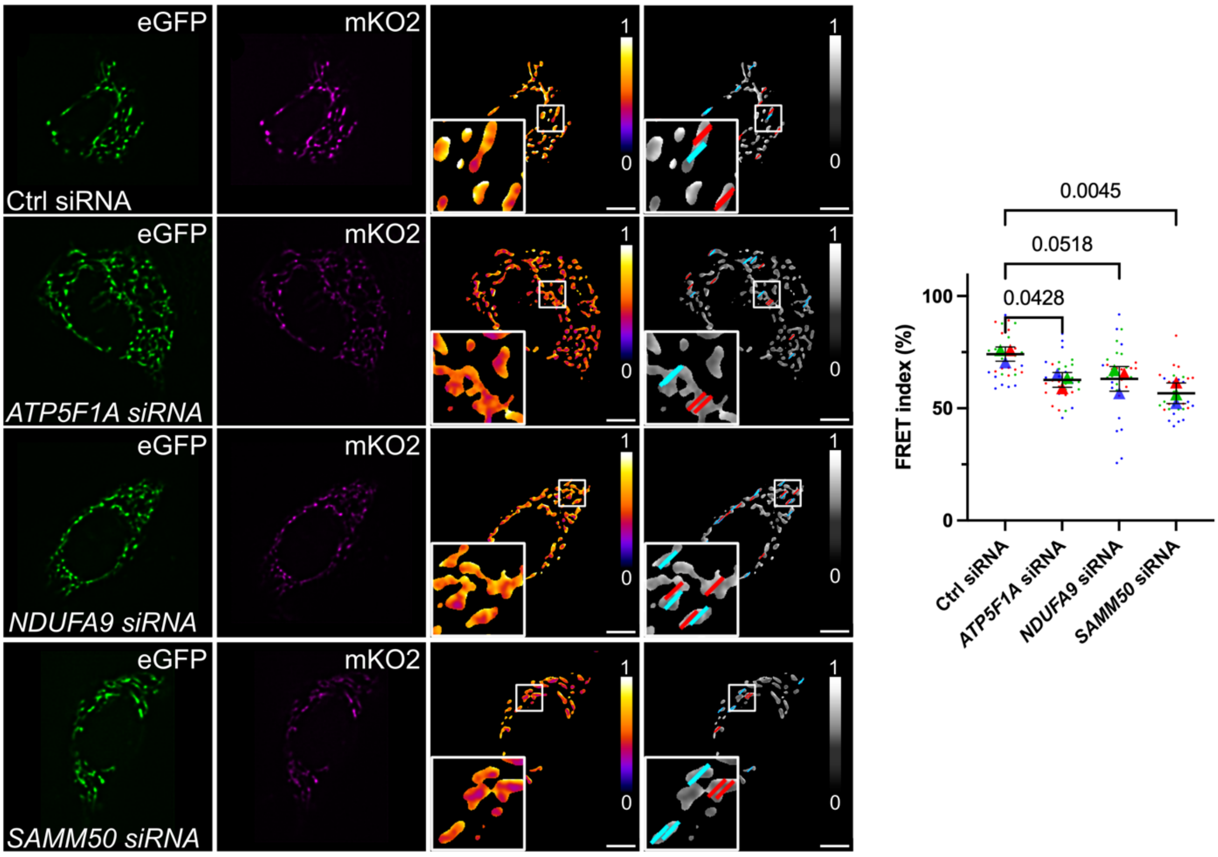


B

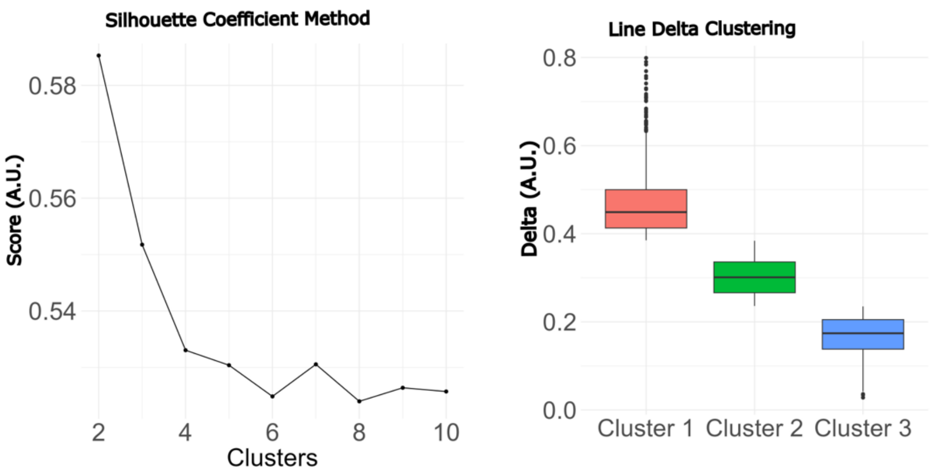


C

| | Repartition (%) | | High Delta - Cluster 1 | | Medium Delta - Cluster 2 | | Low Delta - Cluster 3 | |
|---|---|---|---|---|---|---|---|---|
| **Ctrl siRNA** | Hotspots | 47 | 39,74 | 73 | 16,65 | 78 | 43,62 | 79 |
| | Coldspots | 53 | 28,79 | 49 | 27,55 | 47 | 43,66 | 49 |
| ***NDUFA9* siRNA** | Hotspots | 44 | 56,63 | 75 | 9,13 | 77 | 34,24 | 75 |
| | Coldspots | 56 | 40,67 | 45 | 24,60 | 45 | 34,74 | 47 |
| ***ATP5F1A* siRNA** | Hotspots | 50 | 47,88 | 75 | 12,86 | 76 | 39,26 | 79 |
| | Coldspots | 50 | 36,41 | 49 | 23,34 | 50 | 40,25 | 50 |
| ***SAMM50* siRNA** | Hotspots | 50 | 53,29 | 73 | 9,82 | 76 | 36,89 | 76 |
| | Coldspots | 50 | 34,90 | 47 | 20,83 | 47 | 44,27 | 48 |

**Supplementary Fig. 8. ATP5F1A, NDUFA9 or SAMM50 depletion lower ATP production without affecting its mitochondrial patterning.** (**A**) (From left to right) eSRRFreconstructed images of GFP, mKO2 (FRET), ratiometric FRET images and visual representation of ratiometric FRET shown in greyscale upon random line analyses of T47D cells expressing the mitoGO-ATeam2 biosensor and transfected with control, *ATP5F1A*-, *NDUFA9*-, or *SAMM50*-specific siRNAs. Lines corresponding to FRET Hotspots or Coldspots are pseudocolored in red and cyan, respectively. Insets: higher magnification of the squared area. Graph: mean FRET index values (0-100%) in the indicated conditions. *n* = 11 cells per condition (small dots) in each of three experimental replicates. Large symbols indicate mean values for each biological replicate. Data are means ± S.D. Exact *P*-values are indicated for each comparison. Scale bar: 5 µm. (**B**) (Left) Mean silhouette coefficient score (A.U.) as a function of the number of clusters and (right) distribution of the delta of FRET index values across the three identified clusters. Data go from mean to max, and boxplots show the 1$^{st}$ and 3$^{rd}$ interquartile range and the median delta value. Clusters are ordered by decreasing median delta values, highlighting differences in intensity variation among groups. (**C**) Table showing the overall repartition of FRET Coldspots and Hotspots detected with the MitoGO-ATeam2 biosensor in the indicated transfection condition, together with the quantity of FRET Coldspots and Hotspots depending on the degree of variation of the FRET index (clusters). For each cluster and spot type, the FRET index peak value is shown in red (FRET Hotspots) or cyan (FRET Coldspots).

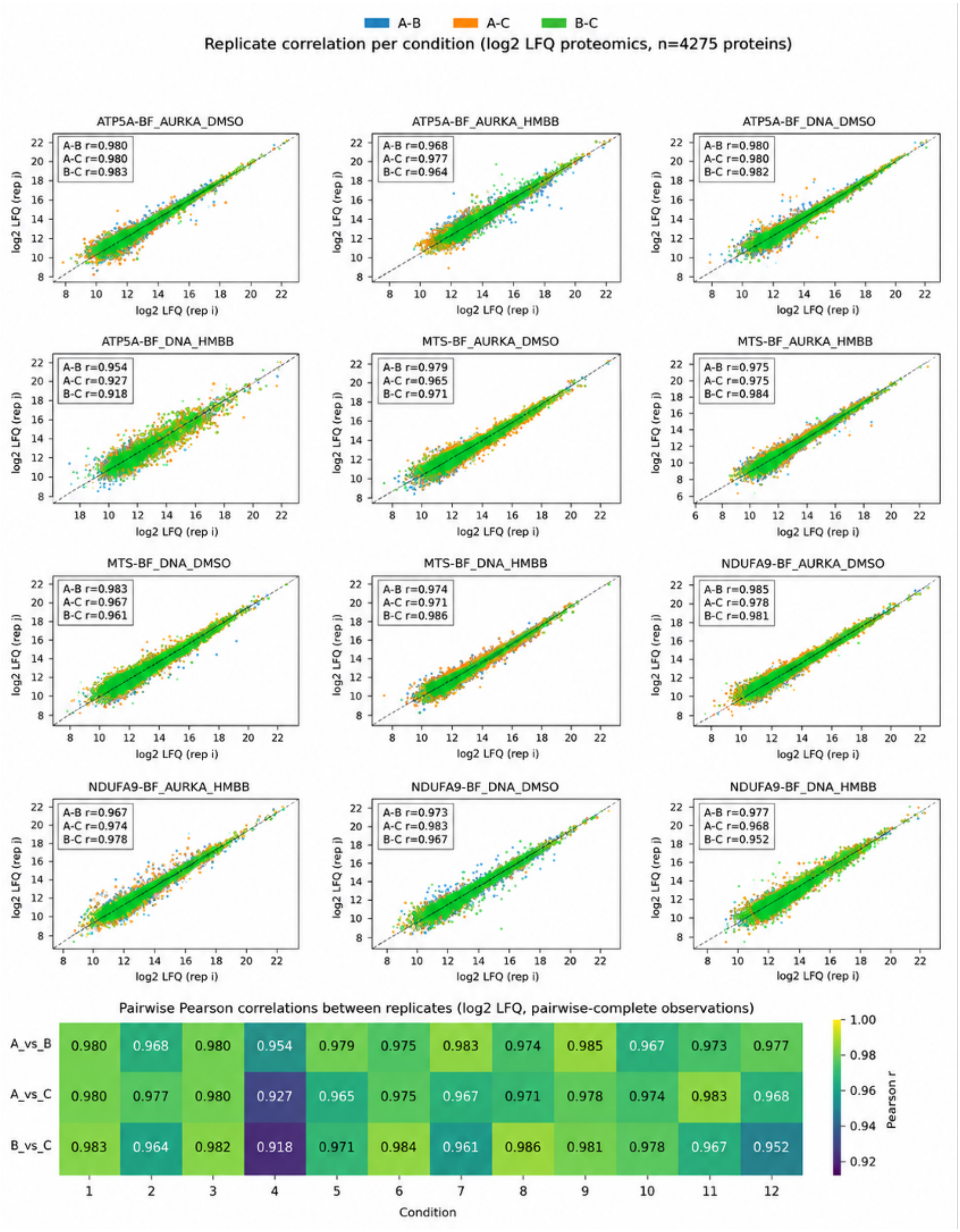


**Supplementary Fig. 9 Correlation analyses between replicates of the TurboID experiments.** Pearson r on log2 LFQ values was calculated for each pair of replicates, as indicated. Conditions are numbered 1-12 in the same order as the panels, from top left (1) to bottom right (12).
